# Probabilistic recovery of cryptic haplotypes from metagenomic data

**DOI:** 10.1101/117838

**Authors:** Samuel M. Nicholls, Wayne Aubrey, Kurt de Grave, Leander Schietgat, Christopher J. Creevey, Amanda Clare

**Affiliations:** Department of Computer Science, Aberystwyth University, Aberystwyth, SY23 3DB, United Kingdom; Department of Computer Science, Katholieke Universiteit Leuven, Celestijnenlaan 200A, 3001 Leuven, Belgium; Institute of Biological, Environmental and Rural Sciences, Aberystwyth University, Aberystwyth, SY23 3DA, United Kingdom; Flanders Make, Oude Diestersebaan 133, 3920 Lommel, Belgium

## Abstract

The cryptic diversity of microbial communities represent an untapped biotechnological resource for biomining, biorefining and synthetic biology. Revealing this information requires the recovery of the exact sequence of DNA bases (or “haplotype”) that constitutes the genes and genomes of every individual present. This is a computationally difficult problem complicated by the requirement for environmental sequencing approaches (metagenomics) due to the resistance of the constituent organisms to culturing *in vitro.*

Haplotypes are identified by their unique combination of DNA variants. However, standard approaches for working with metagenomic data require simplifications that violate assumptions in the process of identifying such variation. Furthermore, current haplotyping methods lack objective mechanisms for choosing between alternative haplotype reconstructions from microbial communities.

To address this, we have developed a novel probabilistic approach for reconstructing haplotypes from complex microbial communities and propose the “metahaplome” as a definition for the set of haplotypes for any particular genomic region of interest within a metagenomic dataset. Implemented in the twin software tools Hansel and Gretel, the algorithm performs incremental probabilistic haplotype recovery using Naive Bayes — an efficient and effective technique.

Our approach is capable of reconstructing the haplotypes with the highest likelihoods from metagenomic datasets without *a priori* knowledge or making assumptions of the distribution or number of variants. Additionally, the algorithm is robust to sequencing and alignment error without altering or discarding observed variation and uses all available evidence from aligned reads. We validate our approach using synthetic metahaplomes constructed from sets of real genes, and demonstrate its capability using metagenomic data from a complex HIV-1 strain mix. The results show that the likelihood framework can allow recovery from microbial communities of cryptic functional isoforms of genes with 100% accuracy.

---

Genomic research is progressing beyond the use of consensus DNA sequences to represent species, towards the ultimate goal of complete characterisation of the genetic diversity that exists across their populations.

So far, research has focused on characterising specific aspects of this diversity, for example: identifying the entire gene-set of all strains of a species (the pangenome) [1]; identifying the groups of genes (or genetic variants within) that are inherited together in organisms across entire populations (the haplome) [2] or in viruses, identifying strains related by mutations in a highly mutagenic environment (the quasispecies) [3].

However many communities (and especially microbial communities) maintain a fine balance between stability and plasticity that is driven both by their genetic breadth and diversity [4, 5, 6, 7]. This necessitates a more holistic approach to allow the simultaneous characterisation of all haplotypes of all organisms in a microbial community (the “metahaplome”).

Complete characterisation of the metahaplome has great biotechnological potential as it would divulge the full repertoire of enzyme isoforms for an organism (or community) and could also guide future antibiotic design [8]. As DNA sequencing technologies advance to produce increasingly longer reads, the depiction of complete haplotypes of species in mixed communities would also become tractable and provide vital missing information for future synthetic biology needs [9].

The general problem of haplotyping, first introduced in 2001 as single individual haplotyping (SIH) [10], has been demonstrated to be computationally difficult (NP-hard) [11] and so focus has moved towards the use of heuristics to make the generation of approximate solutions to SIH both computationally tractable and accurate [12, 13, 14, 15, 16].

However, many current approaches have been developed for diploid species or for those with well-defined genomes such as human [17, 18]. Few are appropriate for organisms where no reference genomes exist or where, in the case of microbes, only a small proportion can be cultured successfully *in vitro.* In these cases, haplotypes must be recovered from DNA isolated and sequenced directly from the environment (metagenomics). Furthermore, in complex datasets such as these, many alternative haplotypes may be possible and there are no methods that can provide the user with an objective function for choosing the best candidates.

The generation of haplotypes from metagenomes is particularly difficult as existing *de novo* analysis pipelines for DNA sequence data generally assume a single individual of origin and, when applied to metagenomic datasets, remove low level variation and produce single consensus sequences [19]. Even specialised metagenomic assemblers [19, 20, 21] do not aim to solve the problem of recovering haplotypes.

Whilst researchers have identified the problem that consensus assembly poses for the downstream analysis of variants [22] and are moving towards alternative assembly approaches, such as graph-based assembly [23], there is still no method for the recovery of individual haplotypes for regions of a metagenome.

To address this need, we have developed a Bayesian framework capable of recovering and ranking haplotypes using evidence of pairs of Single Nucleotide Polymorphisms (SNPs) observed on sequenced reads. While specifically designed to extract haplotypes from metagenomic data of complex microbial communities, the algorithm is general enough to be applied to any analogous haplotyping problem.

We evaluate our approach on metagenomes from both simulated and real sets of genes and identify the haplotypes with the best likelihoods from Influenza and HIV datasets. We demonstrate how, for the first time, the most likely haplotypes can be recovered with high fidelity from complex metagenomic samples enabling the characterisation of the cryptic diversity in microbial communities.

## Results

### The metahaplome

We define the metahaplome as the set of haplotypes for any particular genomic region of interest within a metagenomic data set. A full mathematical definition is available in Supplementary Section 1.

### Hansel and Gretel

We have developed **Hansel**, a data structure designed to efficiently store variation observed across sequenced reads, and **Gretel**, an algorithm that leverages **Hansel** for the recovery of haplotypes from a metagenome. Advantages include that our method:

- recovers haplotypes from metagenomic data
- does not need *a priori* knowledge of the number of haplotypes
- makes no assumptions about the distribution of alleles at any variant site
- does not need to distinguish between sequence error and variation
- uses all available evidence provided by the raw reads
- does not require any user-defined parameters
- does not require bootstrapping, model building or pre-processing
- can confidently rank its own results based on calculated likelihoods
- can be executed on an ordinary computer

The details of the data structure and algorithm are provided in the Online Methods. We provide open source implementations for the data structure API (**Hansel**) and the haplotype recovery algorithm (**Gretel**) at https://github.com/samstudio8/gretel.

We show in *silico* that recovery of haplotypes from simple metahaplomes is possible even with data sets consisting of short reads. Recovery success depends on the availability of sufficient read coverage and enough variation to ensure that all pairs of adjacent SNPs are covered by at least one read. The following subsections describe the datasets used to evaluate **Hansel** and **Gretel**, and the results of this evaluation.

### Synthetic metahaplomes

We evaluated the fidelity of the haplotype reconstructions from **Hansel** and **Gretel** using synthetic metahaplomes. Each synthetic metahaplome consisted of five 3000 bp haplotypes generated by simulated evolution using **seq-gen** [24], with a fixed mutation rate and a star phylogeny (see Methods). Five replicates of seven different mutation rates were generated for a total of 35 metahaplomes.

For each of the 35 metahaplomes simulated by **seq-gen**, we generated 180 sets of uniformly distributed pseudo-reads consisting of 10 replicates for each pairing of 3 read sizes and 6 per-haplotype depths. For the purpose of read alignment and variant calling, we aligned each read set against the 3000 nt starting sequence initially provided to **seq-gen**. Variants were called by assuming any heterogeneous genomic position over the aligned reads was a SNP.

A single run of **Gretel** will repeatedly recover haplotypes until the stopping criteria specified is met (see Methods). For each synthetic metahaplome replicate, we evaluated the fidelity of haplotypes reconstructed by **Gretel** though comparison with the input sequences used to generate the data. The reconstructed haplotype sequence with the greatest proportion of heterozygous positions in agreement with each of the original simulated sequences were determined. We present this recovery rate for the seven mutation rates in combination with the 3 read sizes and 6 per-haplotype read depths used (Fig. 1).

**Figure 1:**
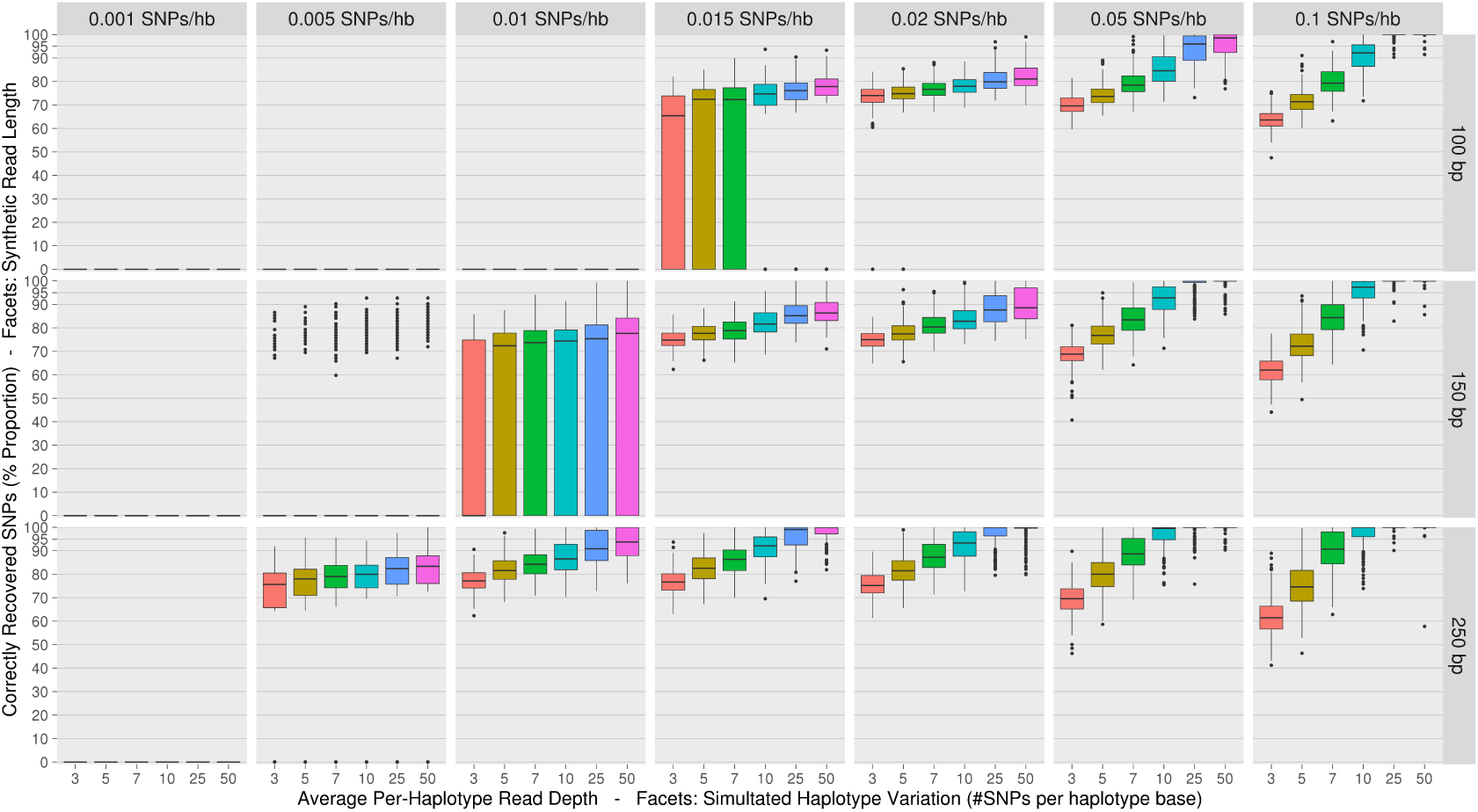
Boxplots summarising the proportion of variants on an input haplotype correctly recovered (y-axes) from groups of synthetic metahaplomes by **Gretel**. Single boxplots present recoveries from a set of five metahaplomes generated with some per-haplotype mutation rate (column facets), over 10 different synthetic read sets with varying read length (row facets) and per-haplotype read depth (colour fill). Each box-with-whiskers summarises the proportion of correctly recovered variants over the 250 best recovered haplotypes (yielded from 50 **Gretel** runs (5 metahaplome replicates × 10 read sets), each returning 5 best outputs). We demonstrate better haplotype recoveries can be achieved with longer reads and more dense coverage, as well as the limitations of recovery on data exhibiting fewer SNPs/hb. This figure may be used as a naive lookup table to assess potential recovery rates for one's own data by estimating the level of variation, with the average read length and per-haplotype depth.

We found that haplotype recovery improves with longer reads and greater coverage. We also observed potential lower bounds on our ability to recover haplotypes from a data set, as the facets with no successful recoveries show. Unsuccessful recoveries are a result of at least one pair of adjacent variants failing to be covered by any read, which is a requirement imposed on **Gretel** for recovery (see Methods). For shorter reads, low-level variation is more of a problem. 0.01 SNPs per haplotype base (hb) over 100 bp would yield just one SNP on average - insufficient evidence for **Gretel**.

Although one might expect high levels of variation to make the recovery of haplotypes more challenging, an abundance of variation actually provides more information for **Gretel**. We observe successful recoveries from data sets with high variation (0.1 SNPs/hb over five haplotypes of 3000 nt yields ≈ 1500 SNPs [Table 4]). With enough coverage (≥ 7x per-haplotype depth), recoveries at a high level of variation are more accurate than those in data sets with fewer SNPs.

For realistic levels of variation (0.01−0.02 SNPs/hb) [25], with per-haplotype read depth of ≥ 7x, we can recover haplotypes at a median accuracy of 80%. With higher per-haplotype depth (≥ 25x), **Gretel** is capable of recovering haplotypes with 100% accuracy (Fig. 1).

### Metahaplomes from real genes

To extend validation to data derived from real genes, and assess the robustness of the likelihood rankings of reconstructed haplotypes, we created a metahaplome consisting of five dihydrofolate reductase (*DHFR*) genes from multiple species (Table 1).

**Table 1:**
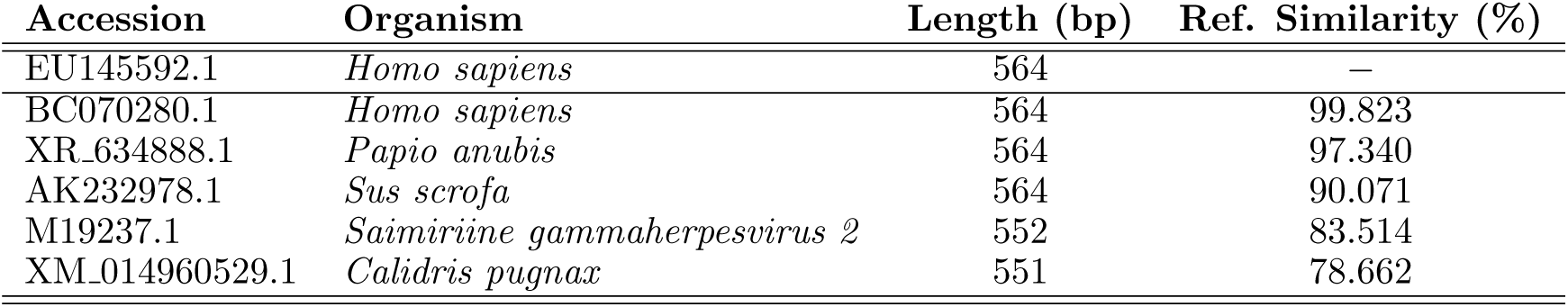
The chosen `pseudo-reference' (EU145592.1) and the five genes that constitute the synthetic DHFR meta-haplome. Genes with decreasing similarity to the reference were selected to pose a more challenging recovery problem. Figure 3 presents our recovery results for each gene.

*DHFR* is an essential enzyme in nucleic and amino acid synthesis and is an important therapeutic target for infectious diseases such as malaria [26]. Identifying *DHFR* haplotypes can help researchers understand how sequence variation contributes to drug resistance.

In a real metagenomic data set, we would use an assembly as a pseudo-reference to align sequenced reads against a genomic region of interest. For simulating the problem of discovering the variants across *DHFR* genes from many organisms in a metagenome, we selected an arbitrary *DHFR* gene (EU145592.1) to serve as the pseudo-reference (Table 1). This reference is not used by **Gretel** for the recovery of haplotypes, but is currently a necessary step to produce an alignment from which to call for SNPs.

Our methodology for testing follows that of the previous section. From the five chosen *DHFR* genes we generated 100 replicates of pseudo-reads for each of the 6 different per-haplotype read depths and 2 read lengths.

The reads were then aligned to the pseudo-reference and the haplotypes recovered using **Gretel**.

Figure 3 presents the proportion of correctly recovered SNPs for each of the five input sequences listed in Table 1. **Gretel** achieves excellent recovery of *BC070280* and *XR_634888* with reads of just 50 bp. Figure 3 also shows that while the accuracy of the reconstruction is influenced by the similarity to the reference used, this can be overcome with longer reads or greater read depth.

**Gretel** scores the haplotypes it recovers with a likelihood, representing the probability of the data observed in the **Hansel** matrix, given the existence of that haplotype. In general, the best reconstructed DHFR haplotypes had the highest likelihoods (Fig. 2). This also improved as the similarity to the reference increased or when there was greater read length or depth.

**Figure 2:**
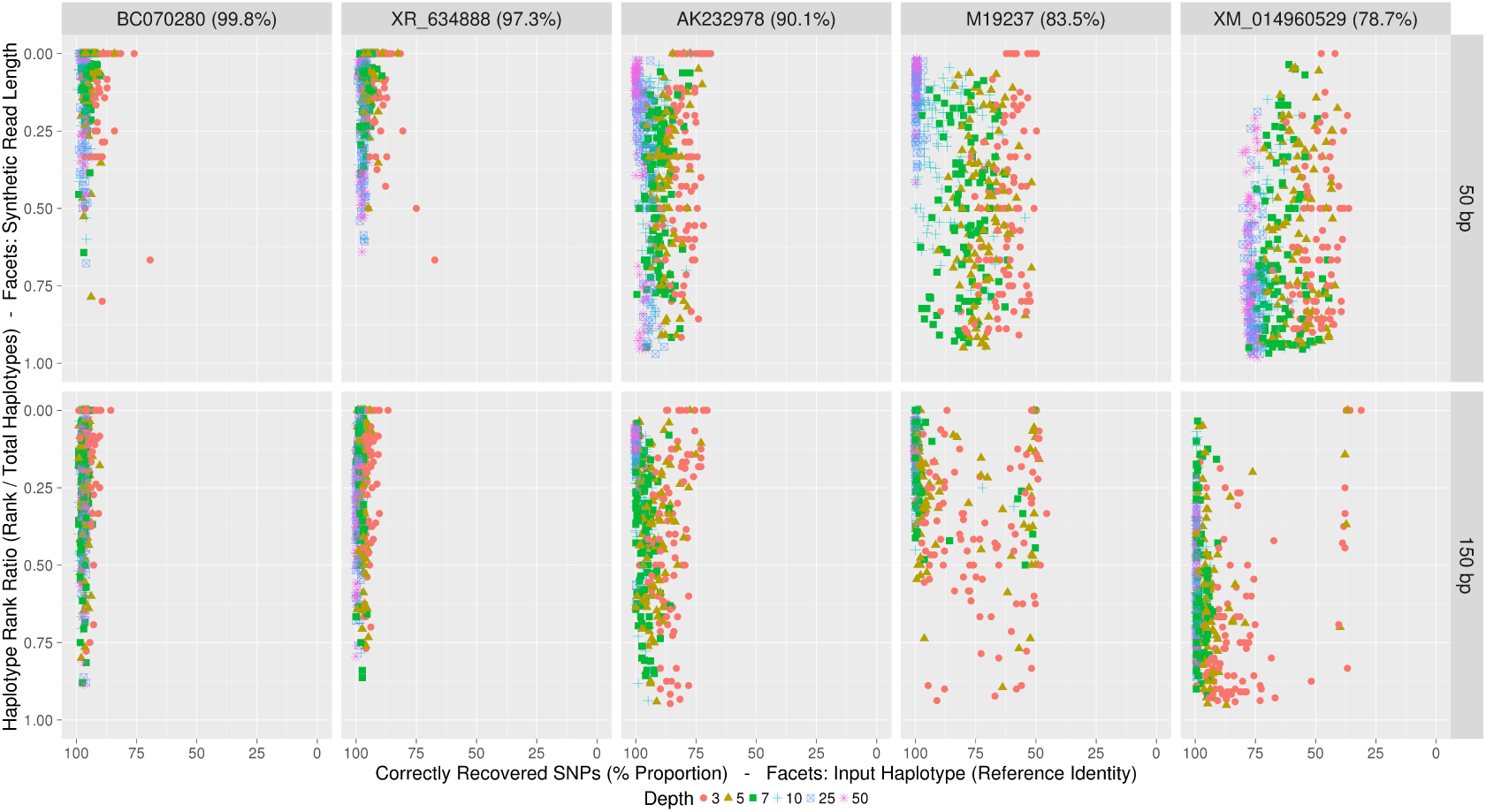
**Gretel** recovery rates (x-axes) against haplotype rank ratio (y-axes) for each gene of the *DHFR* data set (column facets) across the 1200 read sets. Each haplotype recovered has an associated likelihood as scored by **Gretel**. Haplotypes are sorted by their likelihood to obtain rankings. The rank is divided by the total number of recovered haplotypes (for that read set) to produce a rank ratio. A rank ratio of 0 shows a haplotype has the best likelihood awarded by **Gretel**. We present recovery rates against rank ratio, separating the read sets by their read length (row facets) and coverage (colour fill and symbol). **Gretel** is capable of discerning accurate haplotypes, awarding higher rankings to haplotypes with the most identity to real genes.

**Figure 3:**
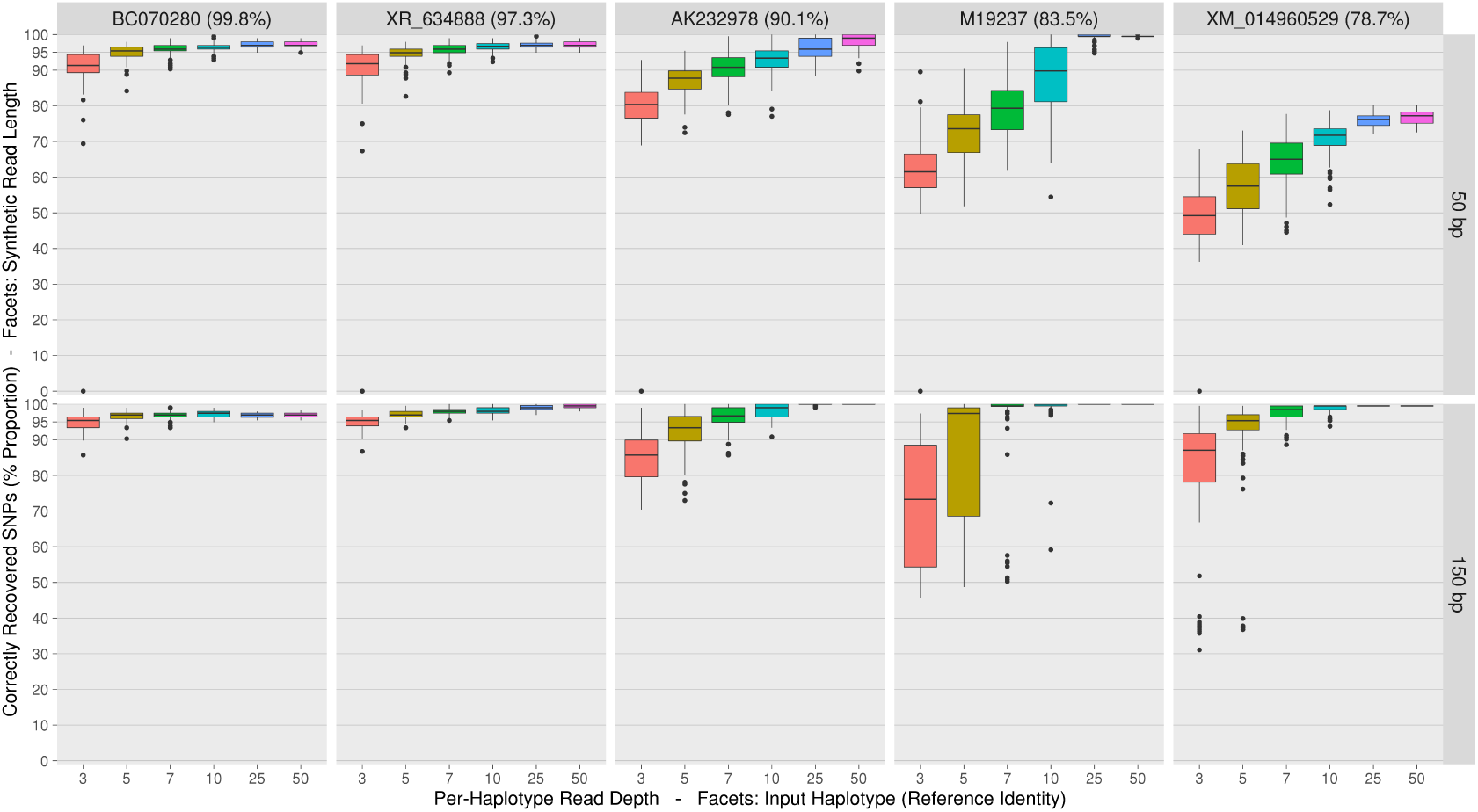
Boxplots summarising the proportion of variants correctly recovered (y-axes) for each input *DHFR* gene (column facets) by **Gretel**. We generated reads from the *DHFR* metahaplome at 6 different per-haplotype read depths (x-axes) between 3 and 50x, 2 read lengths (50 bp and 150 bp row facets) with 100 replicates. Individual box-with-whiskers summarise the recovery rate for a given gene, from reads of a per-haplotype depth and size, over the 100 replicate read sets. Input genes are sorted by decreasing identity to the pseudo-reference left to right. Note that there are a few cases where recovery is not possible at the lowest depth.

We observed that reads from more divergent haplotypes are more readily discarded during alignment to the pseudo-reference, denying **Gretel** the evidence necessary to call these SNPs and hence recover these sequences (Supplementary Fig. 2). Additionally, as some of the haplotypes are more similar to each other, there possibly exists more evidence for the variation shared by those haplotypes, increasing their recovery rate.

Despite this, with sufficient depth and read length, more dissimilar haplotypes can still be recovered with very high accuracy. We observe a similar trend even with increased numbers of input haplotypes (Supplementary Section 11). We encountered an upper limit on the accuracy of recoveries that could not be overcome with additional coverage or increase the read size (Supplementary Fig. 3). This is potentially due to one or more of:

- Shared variation between highly similar haplotypes causing paths in the real metahaplome to converge, making it harder to disentangle the real variation
- Reads discarded during alignment causing pairwise SNP evidence to be withheld from **Gretel**
- Reads discarded during alignment removing evidence for the existence of a heterogeneous site

Despite this, our results show that **Gretel** is capable of recovering haplotypes from synthetic readsets with high accuracy given sufficient read depth, read length or similarity to the reference chosen.

### Metahaplomes from real reads: HIV 5 strain mix

Finally we apply our approach to a set of real reads (*ENA:SRR961514*) consisting of five distinct HIV strains mixed *in vitro* and sequenced on an Illumina MiSeq [27]. Although the sequences of the five strains were known prior to mixing, the strains are likely to have mutated prior to the sequencing of the samples, introducing cryptic diversity [27]. Furthermore, the sequencing process can add additional noise through sequencing error. This provides a challenging data set for haplotype recovery. For the five longest genes on the HIV-1 genome (gag, pol, vif, env and nef) we used **Gretel** to recover all haplotypes present using the real sequencing reads from this experiment.

Table 2 shows, for each gene from each original HIV strain, the similarity of the nearest matching reconstructed haplotype. We found that the *env* gene had the most novel diversity, with the closest matching haplotypes to the original HIV strains only occurring in the top 25 most likely reconstructions. This is in contrast to *pol,* where the closest matching haplotypes to the original HIV strains were in the top 6 most likely reconstructions. This likely represents differing numbers of novel cryptic haplotypes of these genes and correlates well with their known mutation rates *in vivo* [28].

**Table 2:** For each gene from each original HIV strain, the percentage similarity of the nearest matching reconstructed haplotype. Results are presented for each of the five longest genes of the HIV-1 genome (*gag, pol, vif, nef, env*). The bracketed figure indicates the rank of the best haplotype for the strain amongst all recovered haplotypes, according to its likelihood score. We also report the total number of haplotypes recovered by **Gretel** for each gene. *Recovery of 89.6 *env* gene has just one incorrect SNP (2567 SNPs recovered). †Number of haplotypes returned after conservative −1000 *log*_l0_ likelihood cutoff.

| Gene | SNPs | Haplotypes<sup>†</sup> | Strains |  |  |  |  |
| --- | --- | --- | --- | --- | --- | --- | --- |
|  |  |  | 89.6 | HXB2 | JRCSF | NL43 | YU2 |
| *gag* | 1500 | 24 | 100.0 | 100.0 | 99.33 | 100.0 | 99.40 |
|  |  |  | (2) | (6) | (4) | (3) | (10) |
| *pol* | 3009 | 38 | 99.73 | 99.20 | 99.67 | 100.0 | 98.44 |
|  |  |  | (4) | (3) | (2) | (1) | (6) |
| *vif* | 576 | 38 | 100.0 | 98.96 | 100.0 | 100.0 | 97.40 |
|  |  |  | (2) | (9) | (3) | (1) | (5) |
| *nef* | 618 | 60 | 100.0 | 97.14 | 97.30 | 97.62 | 96.02 |
|  |  |  | (2) | (6) | (15) | (5) | (12) |
| *env* | 2568 | 66 | 99.96* | 97.90 | 99.69 | 98.83 | 99.45 |
|  |  |  | (1) | (11) | (7) | (12) | (25) |

For all of the recovered HIV-1 haplotypes, Figure 4 plots the scaled **BLAST** bitscores against the **Gretel** likelihood score. Each recovered haplotype is matched to its closest strain. Our results show that ordering the recovered haplotypes by their likelihood scores can be used as a method to find the best recoveries amongst the recovered sequences.

**Figure 4:**
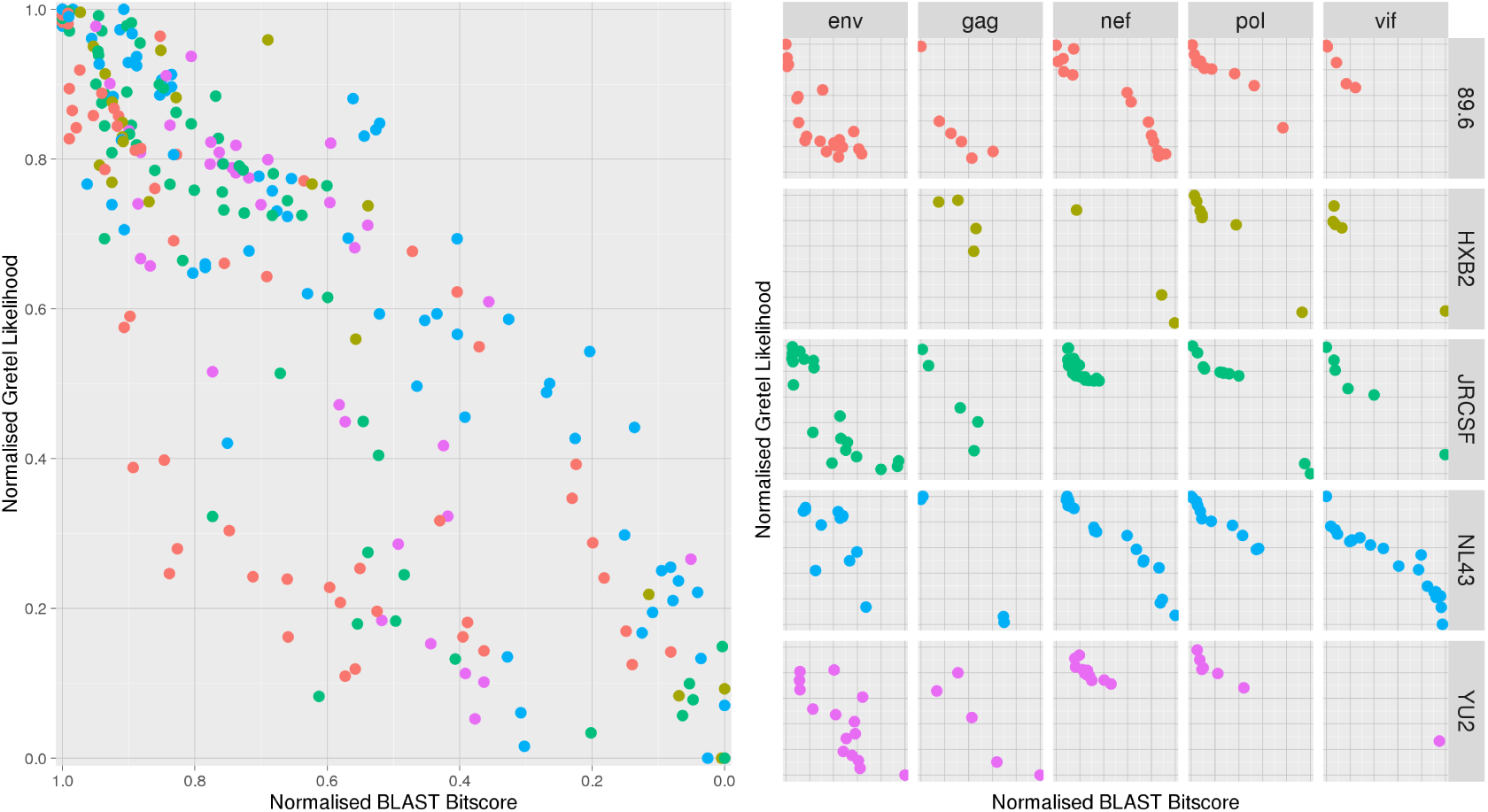
Scaled **BLAST** bitscores (x-axes) against scaled **Gretel** likelihoods (y-axes). (Left) Bitscores against likelihoods for all recovered HIV-1 gene haplotypes, coloured by strain, (Right) Plot facets show the bitscores against likelihood for all recovered haplotypes, separated by gene (column facets) and strain (row facets and colours). **Gretel** consistently awards higher likelihoods to recovered haplotypes that are better matches to a real haplotype.

As reported in Table 2, haplotypes were discarded if they did not meet a conservative threshold of −1000 *log*_10_ likelihood. Manual inspection indicated that these discarded haplotypes showed over selection of deletions, an artifact arising from **Gretel** exhausting non-deleterious evidence in **Hansel** before terminating. Such haplotypes yield no significant **BLAST** hits against the NCBI nr database [29] and our likelihoods provided a clear distinction between noise and useful recoveries.

Beyond the alignment, **Gretel** does not require read processing, parameter bootstrapping or error correction. Despite thousands of heterogeneous sites (primarily caused by sequencing error and alignment noise) **Gretel** is still capable of recovering known haplotypes with 100% accuracy and biologically relevant cryptic haplotypes from metagenomic data.

## Discussion

We offer the term **metahaplome** to represent the set of haplotypes for any particular region of interest within a metagenomic data set. The elucidation of the sequences of the individuals within the metahaplome provides a rich resource of information, enabling detailed study of microbial communities. Synthetic exploitation of the variation observed can be used to improve industrial processes such as biorefining, biomining and synthetic biology [30, 31].

We introduced **Hansel**, a novel data structure which stores the variation observed over reads from a sequenced metagenome. **Hansel** permits traversal of that variation like a graph, yet features probabilistically weighted edges, allowing the current haplotype recovered so far to influence the variants that will be selected next. We also introduced **Gretel**, an algorithm that builds upon **Hansel** for the recovery of genuine haplotypes from a metahaplome constructed from the raw reads of a metagenomic data set.

Together **Hansel** and **Gretel** form a new framework for the recovery of haplotypes in metagenomes, allowing data sets where short read length has previously restricted effective analysis.

### Performance and tractability

We have evaluated **Hansel** and **Gretel** on haplotypes generated by **seq-gen** in order to measure performance with regard to mutation rate, read length and coverage (Fig. 1). We demonstrate very high recovery rates, even in the presence of many SNPs.

We also evaluated our approach with synthetic reads generated from metahaplomes consisting of mixtures of real genes and demonstrated it is possible to recover accurate haplotypes even from short read data (Fig. 3). Successful recoveries can be made with our framework even in the presence of many haplotypes as demonstrated by the high recovery rates from the Flu-A7 metahaplomes, containing short reads generated from 24 diverse influenza sequences (Supplementary Fig. 3).

Finally, we applied our work to a data set consisting of real sequencing reads: the HIV 5 mix metahaplome. We have shown that we can recover long (>2000bp) genes from this complex community with 100% accuracy (Table 2).

Our approach is sensitive to the quality of the alignments of reads against the pseudo-reference, and the choice of pseudo-reference itself. We observed small fluctuations in recovery accuracy across the 24 haplotypes of the Flu-A7 data set depending on the selected reference. During testing of both the DHFR and Flu-A7 data set, we found this sensitivity was because many of the synthetic reads yielded from less similar sequences would not align back to the pseudo-reference (Supplementary Fig. 2), regardless of chosen alignment software.

This does raise an important caveat to our work: both assemblers and aligners will exert influence over the tractability of how many and how accurately haplotypes in a given metagenome can be recovered. Here, the discarding of reads during alignment denies **Gretel** access to critical evidence required to reconstruct those particular haplotypes.

It should be noted that the pseudo-reference is not used by **Hansel** or **Gretel**, it serves only as a common sequence against which to align raw reads. Sequences that happen to share identity with the pseudo-reference are recovered by **Gretel** from the evidence in the **Hansel** matrix, the reference confers no advantage over any other haplotype. Very high recovery rates on sequences that share identity with the pseudo-reference are a reflection of the strength of our approach, and not a trivial recovery. A reference-free method for SNP calling, or a method that constructs or amends the reference from the SNPs [32] would be equally useful to us.

Perhaps most significantly, the tractability of the problem is bound by the quality of the data available. As stated by Lancia [10], it is entirely possible that, even without error, there are scenarios where data is insufficient to successfully recover haplotypes and the problem is rendered impossible.

Our framework has been designed for the recovery of haplotypes from a region of interest in a metagenome (such as variants of a gene involved in a catalytic reaction of interest, e.g. degradation of biomass), but given sufficient coverage of SNPs, our approach could work on regions significantly longer than that of a gene if desired and with data consisting of significantly longer reads.

Regarding time and resource requirements, **Hansel** and **Gretel** is designed to work on all reads from a metagenome that align to some region of interest on the pseudo-reference. Typically these subsets are small (on the order of 10-100K reads) and so our framework can be run on an ordinary desktop in minutes, without significant demands on disk, memory or CPU. Run-times on data with very deep coverage, or many thousands of SNPs, such as the HIV 5-mix, run on the order of hours, but can still be executed on an ordinary desktop computer.

### Methodological comparison of our approach

In contrast to other methods, **Gretel** aims to make as few assumptions as possible. More importantly, our framework requires no configuration, has no parameters and is designed for metagenomic data sets where the number of haplotypes is unknown.

Most SNP calling algorithms discard SNP sites that feature three or more alleles (i.e. non bi-allelic sites) as errors, or under the assumption that input data represents sequenced reads from a diploid species [33]. Although **ParticleHap** [33] relaxes this assumption, it is to reduce the risk of erroneously called genotypes preventing reconstruction of the two haplotypes for a diploid genome.

Many existing methods rely on discarding or altering observed SNPs until a pair of haplotypes can be determined [13]. **Hansel** and **Gretel** use all available pairwise observations and work to recover the most likely haplotypes. Unlike other methods, we do not assume that the observed evidence must be contaminant or sequencing error that needs discarding or altering to recover the real haplotypes. Errors will be poorly supported by read data (and so will have a low probability) and are unlikely to be selected by **Gretel** to become part of a haplotype during traversal.

Most haplotype reconstruction tools are designed specifically for diploid resolution. Some, such as **HapCompass** [18] are designed to identify haplotypes for a single polyploid organism. Such alternatives require prior knowledge of the number of expected haplotypes [34], or other properties of the environment, which are unknowns for metagenomic data sets.

There have been several recent probabilistic approaches targeting the challenge of more complex scenarios such as viral quasispecies assembly [35, 36]. **QuasiRecomb** [37] uses a hidden Markov model solution taking into account recombination events. We tested **QuasiRecomb** on the *gag* region of the HIV dataset, and it predicted over 9000 haplotypes. Optimizing its set of parameters is non-trivial. **PredictHaplo** [38] is also capable of performing full recoveries of the HIV-1 non-**env** genes, but has been shown to recover conservatively outside of HIV-1. [39, 40]. **PredictHaplo** no longer appears to be maintained and we were unable to execute the software on data other than its own test data. The more recent **ViQuaS** [40] reports higher recall than **PredictHaplo**, as expected for an algorithm based on an overlap assembler. However its precision is influenced by the quality of the available reference for the post-assembly filtering step, making **ViQuaS** less suitable for the analysis of a metagenome, where a good reference is unavailable.

With the advent of long-read technology, **ProbHap** recognised a niche in applying computationally expensive dynamic programming solutions to low coverage long-reads [41]. These solutions are inappropriate for high-depth short read data sets as the run time increases exponentially with coverage.

**Lens** [9] is a greedy algorithm for the assembly of long-reads from overlapping short reads, with an algorithm similar to that of **FastHare** [42]. **Lens** uses a straightforward overlap assembly approach that can be used for haplotype resolution as it makes few assumptions. **Lens** generates a large number of unordered haplotypes, including many false positives, but for clean enough datasets this approach may be quick and sufficient. In our tests on the HIV data, **Lens** produced 206 haplotypes for the *env* region and 147 for *pol* without any information on their ranking.

Algorithms that find the correct set of haplotypes from sequence data have the same evaluation issues as information retrieval algorithms. As well as measuring the per-base haplotype similarities, the amount of false positives and false negatives are also important. A brute force algorithm for the production of all possible haplotypes would produce perfect retrieval of the real haplotypes, but would also retrieve many false positives. **Gretel** overcomes this problem by ranking the haplotypes it returns based on the likelihood of that haplotype given the evidence in the **Hansel** matrix at the time that haplotype was completed.

The likelihoods output by **Gretel** are amenable to further statistical analyses such as likelihood ratio testing, or model selection tests using Akaike or Bayesian information criterion [43] to test the fit of complex combinations of haplotypes as descriptions of the sequencing read data.

### Future work

Although we demonstrate **Gretel**’s capability to make perfect recoveries of haplotypes, there exists room for further work. We intend to revisit the following aspects of our approach:

#### • Reweighting

**Gretel** is a deterministic algorithm. The pairwise SNP observation that contributed to the most recent path are reweighted in the **Hansel** matrix to encourage new paths to be discovered. We found that our current choice of reweighting scheme explains some mismatches made by **Gretel** on the *DHFR* and *Flu-A*7 data sets. Other choices of reweighting scheme are possible and we are investigating their potential.

#### • Naive insertion handling

Due to a size constraint on the **Hansel** matrix, further thought is needed to devise a practical methodology that permits the proper consideration of insertions. However, unlike many other approaches **Gretel** does not discard reads containing insertions.

#### • Greediness

**Gretel**’s algorithm involves a greedy bias: We assume the “best” haplotype is the most likely haplotype, and that it can be recovered by selecting the edge with the highest probability at each SNP. However it is possible that **Gretel** could locate solutions whose overall likelihood may be higher with an alternative search strategy.

#### • Stopping Criterion

**Gretel** will generate haplotypes until a dead end in the **Hansel** matrix is encountered, from which there is no evidence for any further transitions. Although we found that our approach can yield low-quality haplotypes before this time, they will have lower likelihoods.

#### • Unused Evidence

There remain sources of evidence not currently used by our algorithm – namely paired end reads and alignment base quality scores. Such data will certainly provide useful co-occurrence and confidence information for SNPs that span some known insert, however careful consideration on how to integrate this data to our approach is necessary.

## Conclusion

In this work we have introduced a definition for the **metahaplome** and provide an implementation of a novel data structure for the storage and manipulation of the evidence supporting the haplotypes within **Hansel**. This tool has value outside of this work, and can provide future algorithms a means to interact with the variation observed in a set of sequenced reads. We also provide **Gretel**, an algorithm for the recovery of metagenomic haplotypes from the **Hansel** data structure. Together they represent a framework for the reconstruction of the haplotypes with the maximum likelihoods from metagenomic datasets without *a priori* knowledge or making assumptions of the distribution or number of variants. Additionally, the algorithm is robust to sequencing and alignment error without altering or discarding observed variation and uses all available evidence from aligned reads.

Existing bioinformatics tools for the processing and analysis of sequencing data make assumptions that are inappropriate for the analysis of haplotypes from metagenomes [44, 19]. Indeed, our own work still relies on alignment to a common reference in order to call SNPs, which can affect recovery rate, as was seen with the synthetic metahaplomes. We would like to investigate methodologies to abandon the requirement for a common reference (that is, in our terminology the assembly or pseudo-reference) and work solely with read data. This would offer opportunities for introducing fewer assumptions and maintaining the integrity of variants observed across metagenomic data before they reach a framework such as **Hansel** and **Gretel**. However, such approaches are not in widespread use or are still under development [23, 45].

In lieu of this, and despite these issues, a metagenomic assembly provides both a convenient proxy for the raw reads and a pseudo-reference against which to align reads and call for SNPs. This does not diminish the ability to accurately recover and rank haplotypes from metahaplomes constructed from both contrived and real data sets. Rather, we demonstrate that our likelihood framework can allow recovery and ranking of haplotypes from microbial communities with 100% accuracy.

**Hansel** and **Gretel** have the potential for the discovery of novel biological insights from microbiomes. This is not a trivial task; many existing *in silico* and *in vitro* techniques such as rational design have struggled to achieve this goal. Applied to metagenomic data from appropriate microbial communities, our approach will reconstruct the cryptic haplotypes within and allow the characterisation of target proteins responsible for catalytic reactions of biotechnological importance.

## Code Availability and Data Access

Our **Hansel** and **Gretel** framework is freely available, open source software available online at https://github.com/samstudio8/hansel and https://github.com/samstudio8/gretel, respectively.

The code used to generate metahaplomes and synthetic reads for both the randomly generated and real gene haplotypes, and the testing data used to evaluate our methods is also available online via https://github.com/samstudio8/gretel-test

## Acknowledgments

CJC was funded by the Biotechnology and Biological Sciences Research Council (BBSRC) Institute Strategic Programme Grant, Rumen Systems Biology, (BB/E/W/10964A01). WA is funded through the Coleg Cymraeg Cenedlaethol Academic staffing scheme. SN is funded via the Aberystwyth University Doctoral Career Development Scholarship and the IBERS Doctoral Programme.

## Author contributions

All authors discussed and defined the theoretical problem and its solutions, SMN wrote the code and documentation and executed experiments, SMN, WA, CJC and AC chose data and designed experiments, all authors analyzed and interpreted the results and wrote the manuscript.

## Methods

### The metahaplome

We have provided a detailed mathematical definition of the metahaplome in Supplementary Section 1.

To enable recovery of a metahaplome from a metagenome with **Gretel** we require:

- *g*, a known DNA region (for example a gene), to be identified by the user
- *c*[*i*: *j*], the region of contig *c* (from an assembly *C*) which has been identified as having similarity to *g*
- *A*_*c*[*i*:*j*]_, the alignments of the set of reads *R* against the contig region *c*[*i*: *j*]
- *S*_*c*[*i*:*j*]_, the genomic positions determined to be SNPs over the region *c*[*i*: *j*]

A metagenomic assembly (which we refer to as a ‘pseudo-reference’, *C*) can be generated by assembling sequenced reads, with an assembler such as **Velvet** [46]. One may identify a gene of interest *g*, on a contig *c* by similarity search or gene prediction. We refer to gene *g* as the *target.* We want to recover the most likely haplotypes of *g* that exist in the metahaplome.

A subset of reads that align to the target region can be determined using a short read alignment tool such as **bowtie2** [47]. Reads that fall outside the target of interest (*i.e*. reads that do not cover any of the genomic positions covered by the target) can be safely discarded: they do not provide relevant evidence to SNPs that appear on the region of interest.

Variation at single nucleotide positions across reads along the target, can then be called with a SNP calling algorithm such as that provided by **samtools** [48] or **GATK** [49]. To avoid loss of information arising from the diploid bias of the majority of SNP callers [33], our methodology aggressively considers any heterogeneous site as a SNP.

The combination of aligned reads, and the locations of single nucleotide variation on those reads can be exploited by **Hansel** and **Gretel** to recover real haplotypes in the metagenome: the **metahaplome**.

### Hansel: A novel data structure

We present **Hansel**, a probabilistically-weighted, graph-inspired, novel data structure. **Hansel** is designed to store the number of observed occurrences of a symbol *α* appearing at some position in space or time *i*, co-occurring with another symbol *β* at another position in space or time *j*. For our approach, we use **Hansel** to store the number of times a SNP *α* at the *i*’th variant some contig *c*, is observed to co-occur (appear on the same read) with a SNP *β* at the *j*’th variant of the same contig. **Hansel** is a four dimensional matrix whose individual elements *H*[*α*, *β*, *i*, *j*] record the number of observations of a co-occurring pair of symbols (*α_i_*, *β_j_*).

### Different from the typical SNP matrix

Our representation differs from the typical SNP matrix model [10] that forms the basis of many of the surveyed approaches. Rather than a matrix of columns representing SNPs and rows representing reads, we discard the concept of a read entirely and aggregate the evidence seen across all reads by genomic position.

At first this structure may appear limited, but the data in *H* can easily be exploited to build other structures. Consider *H*[*α*, *β*,1, 2] for all symbol pairs (*α*, *β*). One may enumerate the available transitions from space or time point 1 to point 2. Extending this to consider *H*[*α*, *β*, *i*, *i* + 1] for all (*α*, *β*) over *i*, one can construct a simple graph *G* of possible transitions between all symbols. In our setting, *G* could represent a graph of transitions observed between SNPs on a genomic sequence, across all reads Figure 5 shows how the Hansel structure records information about SNP pairs, and shows a simple graph constructed from this information.

**Figure 5:**
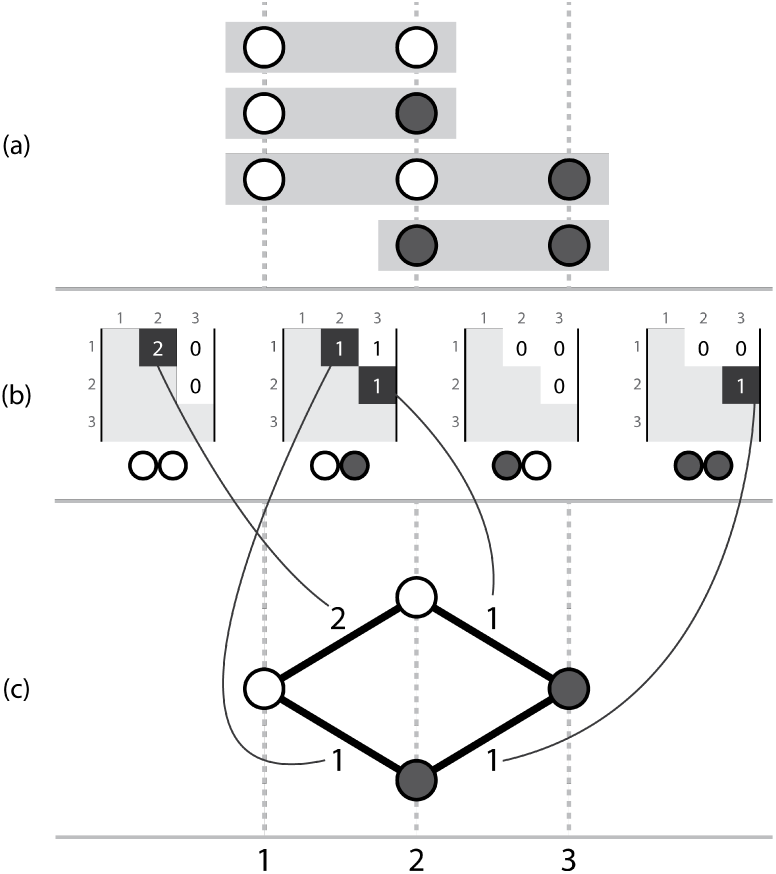
Three corresponding representations, (a) a set of aligned short read sequences, with called variants, (b) the actual **Hansel** structure where each possible pair of symbols (00, 01, 10, 11) has a matrix storing counts of occurrences of that ordered symbol pair between two genomic positions across all of the aligned reads, (c) a simple graph that can be constructed by considering the evidence provided by adjacent variants. Note this representation ignores evidence from non-adjacent pairs, which is overcome by the dynamic edge weighting of the **Hansel** data structure’s interface.

Intuitively, one may traverse a path through *G* by selecting edges with the highest weight in order to recover a series of symbols that represent an ordered sequence of SNPs that constitute a haplotype in the metahaplome. The weight of an edge between two nodes may be defined as the number of reads that provide direct evidence for that pair of SNP values occurring together.

### Different from a graph

Although the analogy to a graph helps us to consider paths through the structure, the available data cannot be fully represented with a graph such as that seen in Figure 5 alone. A graph representation defines a constraint that only considers pairs of adjacent positions (*i*, *i* + 1) over *i*. Edges can only be drawn between adjacent SNPs and their weightings cannot consider the evidence available in *H* between non-adjacent SNP symbols. Without considering information about non-adjacent SNPs, one can traverse *G* to create paths (sequences of SNPs) that do not exist in the observed data set, as shown in Figure 6. To prevent construction of such invalid paths and recover genuine paths more accurately, one should consider evidence observed between non-adjacent symbols when determining which edge to traverse next.

**Figure 6:**
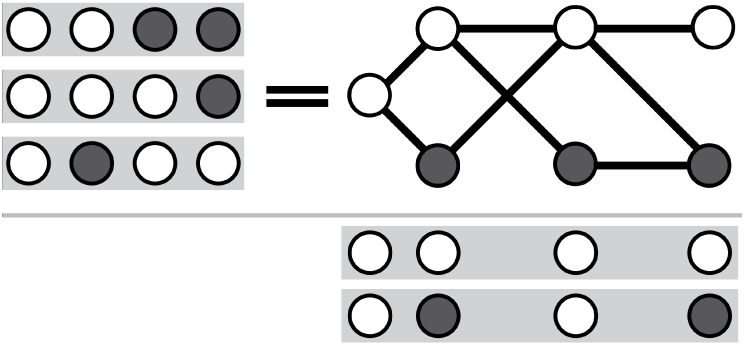
Considering only adjacent SNPs, one may create paths for which there was no actual observed evidence. Here, the reads {0011, 0001, 0100} do not support either of the results {0000, 0101}, but both are valid paths through a graph that permits edges between pairs of adjacent SNPs.

### Using information from non-adjacent SNPs, and the path so far

The **Hansel** structure is designed to store pairwise co-occurrences of all SNPs (not just those that are adjacent), across all reads. We may take advantage of the additional information available in *H* and build upon the graph *G*. Incorporating evidence of non-adjacent SNPs in the formula for edge weights allows decisions during traversal to consider previously visited nodes, as well as merely the current node path, *i*.

That is, given a node *i*, the decision to move to a symbol at *i* + 1 can be informed not only by observations in the reads covering positions (*i*, *i* + 1), but also (*i* − 1, *i* + 1), (*i* − 2, *i* + 1), and so on. Such a scheme allows for the efficient storage of some of the most pertinent information from the reads, and allows edge weights to dynamically change in response to the path as it has been constructed thus far. Outward edges between (*i*, *i* + 1) that would lead to the construction of a path that does not exist in the data can now be influenced by observations in the reads beyond that of the current node and the next. Our method mitigates the risk of constructing paths which do not truly exist.

The consideration and storage of pairwise SNPs fits well with the Naive Bayes model employed to simplify the potentially expensive calculation of conditional probabilities (Supplementary Section 4).

Although we describe **Hansel** as “graph-inspired”, allowing edge weights to depend on the current path through *G* itself leads to several differences between the **Hansel** structure and a weighted directed acyclic graph. Whilst these differences are not necessarily disadvantageous, they do change what we can infer about the structure.

### A dynamic structure

The structure of the graph is effectively unknown in advance. That is, not only are the weights of the edges not known ahead of traversal (as they depend on that traversal), but the entire layout of nodes and edges is also unknown until the graph is explored (although, arguably this would be true of very large simple graphs too). Indeed, this means it is also unknown whether or not the graph can even be successfully traversed.

Also of note is the fact that the graph is dynamically weighted. The current path represents a memory that affects the availability and weights of outgoing edges at each node. Edge weights are calculated probabilistically *during* traversal. They depend on the observation of SNP pairs between some number of the already selected nodes in the path, and any potential next node. Supplementary Section 3 provides the equation and intuition for the probabilistic calculation of edge weights.

In exchange for these minor caveats, we have a data structure that permits graph-like traversal that is intrinsic to our problem definition, whilst utilising informative pairwise SNP information collected from observations on raw metagenomic reads. **Hansel** fuses the advantages of a graph’s simple representation (and its inherent traversability) with the ability to efficiently store pertinent information by considering only pairs of SNPs across all reads.

### Gretel: An algorithm for recovering haplotypes from metagenomes

We introduce **Gretel**, an algorithm designed to interface with the **Hansel** data structure to recover the most likely haplotypes from a metahaplome. To obtain likely haplotypes, **Gretel** traverses the probabilistic graph structure provided by **Hansel**, selecting the most likely SNPs at each possible node (*i.e*. traversing edges with the greatest probability), given some subset of the most recently selected nodes in the path so far. At each node, an *L*’th order Markov chain model is employed to predict which of the possible variants for the next SNP is most likely, given the last *L* variants in the current path.

Execution of **Gretel** can be broken into the following steps:

1. Parse the read alignments and retain only the bases that cover SNP sites, discarding any conserved base positions as they provide no haplotype information.
2. Populate the **Hansel** structure with all pairwise observations from each of the reads.
3. Exploit the **Hansel** graph API to incrementally recover a path until a variant has been selected at each SNP position:

- Query for the available transitions from the current position in the graph to the next SNP
- Calculate the probabilities of each of the potential next variants appearing in the path given the last L variants
- Append the most likely variant to the path and traverse the edge
4. Report this path as a haplotype and then remove the information for this path from the data by reweighting observations that contributed to this path. This will allow for new paths to be retrieved next.
5. Repeat (3−4) until the graph can no longer be traversed or an optional additional stopping criterion has been reached.

### Greedy path construction

Haplotypes are reconstructed as a path through the **Hansel** structure, one SNP at a time, linearly, from the beginning of the sequence. At each SNP position, the **Hansel** structure is queried for the variants that were observed on the raw reads at the next position. **Hansel** also calculates the conditional probabilities of each of those variants appearing as the next SNP in the sequence, using a Markov chain of order L that makes its predictions given the current state of the observations in the **Hansel** matrix and the last *L* selected SNPs. **Gretel**’s approach is greedy: we only consider the probabilities of the next variant. Our razor is to assume that the best haplotypes are those that can be constructed by selecting the most likely edges at every opportunity.

### Reweighting to find multiple haplotypes

Whilst our framework is probabilistic, it is not stochastic. Given the same **Hansel** structure and operating parameters, **Gretel** will behave deterministically and return the same set of haplotypes every time. However, we are interested in recovering the metahaplome of multiple, real haplotypes from the set of reads, not just one haplotype. **Hansel** exposes a function in its interface for the reweighting of observations. Once a path through the graph is completed (a variant has been chosen for all SNP sites), the observations in the **Hansel** matrix are reweighted by **Gretel**.

Currently, **Gretel** reduces the weight of each pairwise observation that forms a component of a completed path - in an attempt to reduce evidence for that haplotype existing in the metahaplome at all, allowing evidence for other haplotypes to now direct the probabilistic search strategy.

### Gretel's outputs

Finally, **Gretel** outputs recovered sequences as FASTA, requiring no special parsing of results to be able to conduct further analyses. Of course, with knowledge of the input haplotypes that we expect to recover, we are able to quantify our approach. For real metahaplomes, we need a mechanism to differentiate false positives, or to rank our confidence in the returned haplotypes.

Future work will investigate this in more depth, but currently, in addition to the sequences themselves, **Gretel** outputs a ‘crumbs’ file — a whimsical name for a simple, tab delimited format — that contains metadata for each of the recovered sequences: log probability of that sequence existing given the evidence seen overall, how much of the evidence in **Hansel** that particular sequence was supported by, and how much of that evidence was reweighted as a result of that path being chosen.

Currently, **Gretel** will continuously recover paths out of the remaining evidence until it encounters a node from which there is no evidence that can inform the next decision.

### Testing methodologies

In this section, we provide an overview of the methods to generate metahaplomes for both synthetic haplotypes, and haplotypes based on real genes. We describe our approach for evaluation of our work. Our test methodologies evaluate the performance of our framework against metahaplomes consisting of synthetic reads derived from both randomly generated haplotypes, and also haplotypes created from real gene sequences. Table 3 summarises each of the evaluation data sets.

**Table 3:** Properties of the evaluation data sets. ^†^Per haplotype base variation rate estimated by dividing the ratio of SNPs over sequence length by the number of known haplotypes. ^‡^Read size and depth for HIV-1 data sets were averaged over the region of the particular gene in the aligned BAM (Table 5).

| Dataset Name | Length (bp) | Variation (SNPs/hb) | Read Sizes (bp) | Per-haplotype Depths | Replicates | Number of Read Sets | Number of Haplotypes |
| --- | --- | --- | --- | --- | --- | --- | --- |
| seq-gen | 3000 | 0.001 | 100, 150, 250 | 3, 5, 7, 10, 25, 50 | 10 × 5 trees | 180 × 5 | 5 |
|  |  | 0.005 |  |  |  | 180 × 5 |  |
|  |  | 0.01 |  |  |  | 180 × 5 |  |
|  |  | 0.015 |  |  |  | 180 × 5 |  |
|  |  | 0.02 |  |  |  | 180 × 5 |  |
|  |  | 0.05 |  |  |  | 180 × 5 |  |
|  |  | 0.1 |  |  |  | 180 × 5 |  |
| DHFR | 564 | 0.0696<sup>†</sup> | 50, 150 | 3, 5, 7, 10, 25, 50 | 100 | 1,200 | 5 |
| Flu-A7 | 982 | 0.01175<sup>†</sup> | 50, 150 | 3, 5, 7, 10, 25, 50 | 100 | 1,200 | 24 |
| HIV-gag | 1500 | 0.2<sup>†</sup> | 219<sup>‡</sup> | 6465.07<sup>‡</sup> | 1 | 1 | 5 |
| HIV-pol | 3009 |  | 217<sup>‡</sup> | 9396.89<sup>‡</sup> |  | 1 |  |
| HIV-nef | 618 |  | 218<sup>‡</sup> | 4211.87<sup>‡</sup> |  | 1 |  |
| HIV-vif | 576 |  | 216<sup>‡</sup> | 4822.18<sup>‡</sup> |  | 1 |  |
| HIV-env | 2568 |  | 214<sup>‡</sup> | 4539.87<sup>‡</sup> |  | 1 |  |

### Read generation and variant calling

With the exception of the HIV data set, our reads are generated *in silico* with our Python based tool (**shredder**) which can be found as part of our evaluation repository via: https://github.com/SamStudio8/gretel-test.

Our synthetic reads are designed to be simplistic; errorless and of uniform length and coverage. The synthetic read sets form a basis for testing the **Hansel** and **Gretel** packages during development, as well as providing a platform on which to investigate the influence that parameters such as read length, number of haplotypes, and mutation rate have on haplotype recovery.

For a given FASTA file, our tool generates reads of a uniform user-defined length and coverage, for each of the sequences in the file. The tool calculates the number of reads to generate to achieve the approximate coverage, given the length of the sequence, and the selected read length. A BED file can be used to mask particular areas of one or more of the input FASTA sequences.

Uniform coverage is approximated by randomly generating the start positions of all of the reads across the input sequence (and also allowing for up to half of a read to fall off either end of the sequence).

As our tool is aware of the start position of every read that it generates, it is possible to also produce an alignment of those reads in SAM format. This allows us to align reads without introducing biases and assumptions from external tools. Pileups of our generated reads typically feature many tri- or tetra-allelic sites (especially as mutation rate increases). Many variant calling approaches feature diploid biases and can disregard such sites as sequencing error, denying us the information with which to recover correct haplotypes. Our approach is robust to noise arising from sequencing error (see the results for the HIV data, which consists of real reads from a mixed set of 5 HIV strains). It is also robust to misaligned reads (see results for the DHFR, Influenza and HIV data) and as such we can aggressively call variants by assuming any heterogeneous site is a SNP.

Our evaluation repository contains the simple **snpper** tool that generates a VCF for a given BAM. **snpper** outputs a VCF record for any heterogeneous site.

The code, documentation, and data for evaluation are open source and freely available via our data and testing repository: https://github.com/samstudio8/gretel-test

### Evaluating recovery accuracy

To evaluate the accuracy of a run of **Gretel**, each known input haplotype is compared pairwise to each of the recovered output haplotypes. Each input haplotype is matched to a corresponding “best” recovered haplotype. Best is defined as the output haplotype that yields the smallest Hamming distance from a given input haplotype. For each synthetic metahaplome, we perform a multiple sequence alignment with **MUSCLE** [50] to determine the definitive SNP positions. When calculating Hamming distance, we consider only these corresponding positions. That is, we exclude the comparison of homogeneous sites from the evaluation metric, to ensure we only consider our accuracy on positions that require recovery. For our results we report the proportion of total SNPs that were correctly recovered by **Gretel**, expressed as a percentage.

Comparing sites enumerated by the multiple sequence alignment of the original haplotypes, as opposed to the VCF of each individual read set ensures **Gretel** is penalised when a SNP has not been called from the read set.

Note that regardless of quality, all input haplotypes are assigned a best output haplotype. An output haplotype may be the best haplotype for more than one input. If more than one output haplotype has the same Hamming distance, the first that was found is chosen. If **Gretel** could not complete at least one haplotype (*i.e.* a pair of adjacent SNP positions were not covered by at least one read), all input haplotypes are awarded a recovery of 0%.

### Synthetic (seq-gen) metahaplomes

With the desire to first test our approach on data sets with well-defined and controllable read properties, but still posing a recovery problem, **seq-gen** [24] was used to generate sets of DNA sequences that would serve as haplotypes of a synthetic metahaplome.

**seq-gen** simulates the evolution of a nucleotide sequence along a given phylogeny. For testing **Gretel**, we provided a star shaped guide tree with uniform branch lengths, such that all haplotypes would be equally dissimilar to each other. These uniform branch lengths correspond to the rate of per haplotype base (hb) nucleotide heterogeneity. Thus, each taxa in the tree has a DNA sequence based upon the evolution of the given starting sequence, following simulated evolution at the given rate.

The same starting sequence was shared by all of our generated trees. We used a randomly generated sequence of 3000 nt with 50% GC content. We fixed the number of taxa in the trees at five, but varied the mutation rate across seven levels (Table 4. 35 trees were generated (7 mutation rates and 5 replicates), each containing five sequences mutated at the same rate, from the original 3000 nt sequence. Each of the resulting 35 sets of five mutated DNA sequences represent a metahaplome from which the five haplotypes must be recovered by **Gretel**.

**Table 4:** Mean number of variants called over the 900 generated synthetic read sets, for each per-haplotype base (hb) mutation rate. The generated sequences for each metahaplome were 3000 nt long.

| Mutation Rate<br>(SNPs/hb) | Average number of<br>variants called |
| --- | --- |
| 0.001 | 17.25 |
| 0.005 | 71.20 |
| 0.010 | 141.54 |
| 0.015 | 209.25 |
| 0.020 | 277.28 |
| 0.050 | 640.63 |
| 0.100 | 1159.62 |

As per our described read generation and variant calling protocol, we generated synthetic reads from each of the five sequences in the metahaplome, varying both the read length and per-haplotype read depth *(i.e.* the average coverage of each haplotype). For each read length and depth parameter pair, ten read sets were generated, to amortise any effect on haplotype recovery introduced by the alignments of the reads themselves. We generated 6300 read sets (3 read sizes, 6 per-haplotype depth levels, 7 mutation rates, 10 read replicates, 5 tree replicates). Table 3 summarises the data sets generated for evaluation.

### Metahaplomes from real genes

~~~
bowtie2 --local
-D 20 -R 3 -L 3 -N 1 -p 8
--gbar 1 --mp 3
-x master.bti -U reads.fq --un unaligned.fq --no-unal -S out.sam
~~~

#### Listing 1 Bowtie2 parameters used to align the read sets

We chose an arbitrary *DHFR* gene from GenBank (*EU145592.1*) to serve as the ‘master’ (pseudo-reference) sequence against which to align reads to call variants of a synthetic *DHFR* metahaplome. To find sequences to use as haplotypes, a discontinuous **megaBLAST** was conducted against the master. A set of five related but arbitrary genes of decreasing sequence identity (≈99.8%, 97.3%, 90.1%, 83.5%, 78.7%) were selected (Table 1).

As per our previously described read generation method, we created read sets generated from the five input sequences, varying their length and depth. Our data set is outlined in Table 3, and consists of 1,200 sets of reads (2 read sizes, 6 per-haplotype coverage levels, 100 replicates).

Unlike the methodology for the **seq-gen** simulations, our *DHFR* reads were aligned back to the pseudo-reference with **bowtie2**, as the input sequences were not of the same length and could potentially contain insertions or deletions with respect to the chosen reference. Reads from the more dissimilar sequences were discarded by **bowtie2** (Supplementary Materials Fig. 2), which is a problem, because the variant information that they contain is then no longer available to add to **Hansel**. This was also true of other commonly used sequence aligners, including **bwa**. These tools are not designed for use-cases where one wishes to permit such diverse sequences to be co-aligned. Though, we were able to improve overall alignment scores for our *DHFR* and Flu-A7 data sets by significantly relaxing **bowtie2**’s parameters (Listing 1).

Variants were called on the alignment using our **snpper** tool (provided at https://github.com/samstudio8/gretel-test) that determined all heterogeneous sites as variants. A multiple sequence alignment of the original five haplotypes showed the number of actual SNPs in the data set was 196. Supplementary Figure 1 shows the proportion of SNPs that were correctly called from the read sets (before recovery). In this scenario, very short reads (50 bp) appear to offer insufficient evidence to call all the real variants. This can prevent full recovery of some of the haplotypes (whose variant sites were lost to synthetic read generation, or alignment).

### Metahaplomes from real reads

For verification of our approach on real sequenced reads, we evaluated **Gretel** with a dataset designed specifically for the benchmarking of haplotype reconstruction methods [27]. Five well studied HIV-1 strains (*89.6, HXB2, JRCSF, NL43,* and *YU2*) were mixed and sequenced with an Illumina MiSeq.

The Illumina reads are available from ENA (*SRR961514*). As per our protocol, reads were aligned against a pseudo-reference. We selected one of the five strains to serve as this reference (*89.6,* and aligned all sequenced reads against it using **bowtie2**. The overall alignment rate was 96.87%, yielding an alignment of 1,385,162 sequences.

We determined that any heterozygous pileup site in the resulting alignment would be defined as a SNP, resulting in a VCF containing 9,570 called variants. The SNPs are so numerous that they occur at 98.98% of all sites.

We executed **Gretel** on five of the largest genes on the HIV-1 genome, using *HXB2* gene co-ordinates [51], each time providing the same alignment BAM and VCF to **Gretel**, but additionally using the **start** and **end** command line parameters to define the boundary of the particular region of interest. Table 5 describes the five genes and their properties. Table 6 shows the identity matrix for the five strains.

**Table 5:** The five HIV-1 genes, HXB2 co-ordinates [51] and associated properties of the aligned reads

| Gene | Region (Size) | Average Coverage | Called SNPs |
| --- | --- | --- | --- |
| *gag* | 790—2289 (1,500 bp) | 32,325.34 | 1,500 |
| *pol* | 2084—5093 (3,009 bp) | 46,984.47 | 3,009 |
| *vif* | 5040—5616 (576 bp) | 24,110.89 | 576 |
| *nef* | 8796—9414 (618 bp) | 21,059.34 | 618 |
| *env* | 6224—8792 (2,568 bp) | 22,699.35 | 2,568 |

**Table 6:** Percentage identity matrix for the five HIV-1 strains as reported by **MUSCLE** [50]

| Strain | *89.6* | *JRCSF* | *YU2* | *HXB2* | *NL43* |
| --- | --- | --- | --- | --- | --- |
| *89.6* | 100.00 | — | — | — | — |
| *JRCSF* | 93.98 | 100.00 | — | — | — |
| *YU2* | 94.05 | 95.10 | 100.00 | — | — |
| *HXB2* | 94.43 | 95.18 | 95.71 | 100.00 | — |
| *NL43* | 94.07 | 94.98 | 95.39 | 97.49 | 100.00 |

We evaluated our approach using the same framework as our synthetic metahaplomes. We report the proportion of variants correctly recovered by **Gretel** for each of the five known haplotypes, for each of the genes.

## Supplementary Information

### 1 The metahaplome

Consider:

- Ω An environment of microbial organisms.
- *O* A bag containing each full genomic sequence, of each individual organism in environment Ω. *O* is the whole metagenome of Ω.
- *o*[*i*: *j*] A sub-sequence *i*..*j* of some genome *o* ∈ *O*
- Gene *g*, Protein *p* A known DNA sequence *g*, responsible for the production of a protein *p*. *p* is a protein capable of performing a catalytic reaction of interest, such as the hydrolysis of cellulose
- Δ(*s g*) Any function Δ that can determine whether a DNA sequence *s* (such as a sub-sequence *o_n_*[*i*: *j*]) has sufficient similarity to *g*. Δ may return a boolean, or a real value with some user-selected threshold.
- Γ*g*=*se*t(⎱*o_k_*[*i*:*j*]|Δ(*o_k_*[*i*,*j*],*g*),*k*∈1..|*O*|,*i*,*j*∈1..|*o_k_*|₰)
- One may collect a bag of sub-sequences found to be a similar to *g* and thus responsible for the production of a protein product *p*. The set Γ*_g_* of unique sequences contained in the bag enumerate the possible sub-sequences across all *o* ∈*O* that are capable of producing a protein product *p*.

We define Γ_*g*_ as the **metahaplome** for the gene *g*, in the metagenome *O*. Each such member γ ∈ Γ_*g*_ is a *haplotype* of *g*, capable of constructing a protein that performs a biological function of interest in the environment. We wish to recover the set Γ_*g*_. Additionally, our methodology requires the following definitions:

- *σ* = Sample(Ω)
  - A sample taken from microbial environment Ω.
  - *σ* ⊂ Ω
- *M*
  - A bag containing each full genomic sequence *m*, of each individual organism captured in the subset σ.
  - *M* represents the metagenome that was captured in the sample *σ*.
  - *M* is not necessarily representative of the entire metagenome *O*.
  - *M* ⊂ *O*
- *R* = Seq(*σ*)
  - A set of sequenced **reads** *r* ∈ *R*, consisting of a series of residues (bases) *r*[*i*] ∈ {*A*, *C*, *G*, *T*, *N*}, *i* ∈ 1..|*r*|.
  - *R* is the set of reads obtained from the sequencing of isolated DNA from sample *σ*.
  - *R* contains fragments of genomes *m* ∈ *M* (and error).
  - Due to sampling bias acquiring *σ* from Ω, *R* is unlikely to be representative of the true genetic diversity available in *O*.
  - Additionally, due to sequencing and PCR biases and error, *R* is unlikely to provide uniform non-zero coverage of the residues across all *m* ∈ *M*.
- *C* = Assembly(*R*)
  - Contig set *C* (the “**Assembly**”) constructed from the reads *R* by some **Assembly** operation.
  - *C* poses as a pseudo-reference for the sequenced metagenome.
  - *C* attempts to estimate *M*, but typically fails to distinguish between similar sequences that belong on distinct *c*.
  - *c*[*j*] ∈ {*A*,*C*,*G*,*T*,*N*} for *j* ∈ 1..|*c*|, for *c* ∈ *C*.
- *A* = Align(*R*, *C*)
  - **Alignment** *A*, the set of alignments constructed via aligning read set *R* to contig set *C* with operation **Align**.
  - *A*_*c*[*i*:*j*]_, the subset of A containing any alignment on contig *c* that spans any site between *c*[*i*..*j*]
- *S* = Call(*A_c_*)
  - The set of genomic positions on a contig *c* ∈ *C* determined to be a SNP (single nucleotide polymorphism) by an operation **Call**, given the subset *A_c_*: the alignments of *R* against *c*.

Our goal is to determine Γ_*g*_, for some gene *g*. Although M will not be entirely representative of *O*, it is the only evidence of the sequence diversity available. Thus we adjust our goal to instead find the most likely elements of Γ_*g*_, given the evidence that can be derived from *M*, via the alignments *A* and variant sites *S*.

### 2 Hansel as a graph

Consider an alphabet of symbols, Σ (*e.g*. {*A,C,G,T, N,* −}) and a list of n SNP positions 1..*n*. Symbols ∅_*S*_ and ∅_*E*_ represent special sentinal positions at the start and end of the SNP positions (0 and *n* +1 respectively). As described in our article, the **Hansel** structure *H* can be considered as a graph *G* = (*V*, *E*). Here, we define *V*, and *E*:

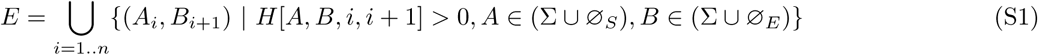

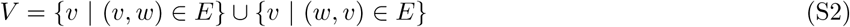

*E* represents the set of edges, where an edge (*A*_i_, *B_i_*_+1_) is determined to exist in *E* if there exists at least one read whereby symbol *A* was observed at position *i* to co-occur with symbol *B* at SNP position *i* +1.

It should be noted, that although *G* can be constructed from *H* such that it is undirected and contains cycles, both properties lead to nonsensical haplotypes. Under such circumstances, **Gretel** could construct a path that visits multiple nodes that appear at the same *i*, or a trail that visits the same node multiple times. Such sequences would be meaningless in the context of haplotype construction, thus the interface to **Hansel** acts in such a way that *G* is a directed, acyclic graph.

We can define a haplotype as an alternating sequence of nodes (*v* ∈ *V*) and edges (*e* ∈ *E*). A path must always start and end at the special sentinel symbols ∅_*S*_ and ∅_*E*_, respectively.

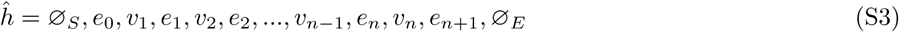

Although, as only one directed edge between some *v_i_* and *v_i_*_+1_ may exist, we can define *ĥ* as a sequence of *v* ∈ *V*:

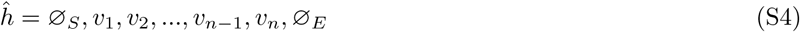

### 3 Probabilistic edge weights

However, if the construction of *G* does not consider elements in *H*[*A*, *B*, *i,j*] where *abs*(*i − j*) > 1 (non-adjacent SNPs) it is likely one will recover haplotypes that do not actually exist.

Given the pairwise information available in *H*, for both adjacent, and non-adjacent SNPs, across all reads, edges in the graph *G* derived from *H* can be weighted probabilistically. We attempt to determine the next most likely symbol in a sequence, considering both the marginal distribution of symbols at the next position and the likelihood of those symbols appearing next, given an already observed partial sequence. That is, the next symbol *v_i_*_+1_ in a path depends not only on the current symbol (*v_i_*) but some number of previous symbols (*v_i_*_−1_, *v_i_*_−2_…*v*_0_).

The outgoing edges from *v_i_* are probabilistically weighted by exploiting the observations stored in the **Hansel** structure to create probabilities. These probabilities then determine the likelihood of moving from some *vi* to each of the possible *v_i_*_+1_.

We take a Bayesian approach to the problem of probabilistically weighting edges in **Hansel**’s graph representation. We define the probability of selecting *v_i_*_+1_, conditioned on the path observed so far:

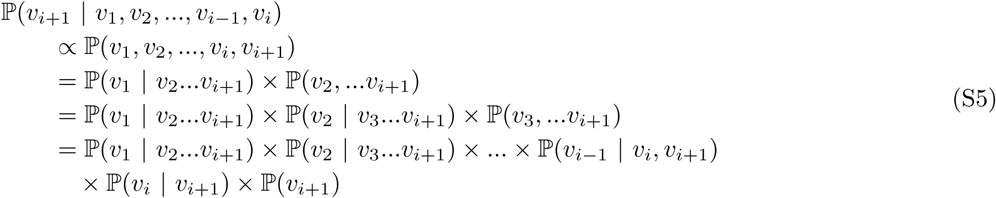

### 4 Simplification of conditional edge weights

Clearly, the number of factors in Equation S5 increases with *i*. For longer paths (more single nucleotide polymorphisms detected along the target region of interest), evaluating the equation becomes more computationally expensive, and risks potentially compounding estimation errors.

To construct a whole path *p* from *v*_1_ …*v*_n_, the upper bound for the number of iterations will be |Σ| × *n* with calculations becoming increasingly complex as *i* increases.

To reduce complexity, we make an assumption of conditional independence between variants. Whilst this seems counter intuitive, the Naive Bayes model can deliver robust results despite its coarse assumption.

Thus we may simplify our previous equation and consider only the pairwise appearances of each *v_i_* encountered thus far against *v_i_*_+1_.

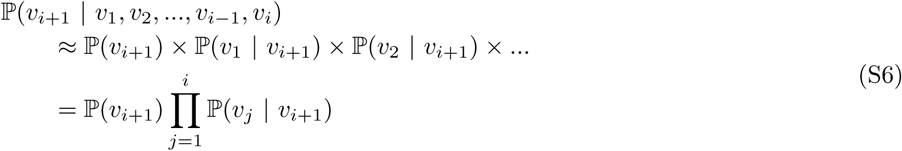

However as discussed in our article, individual reads will not cover all SNP positions 1..*n* (if they did, we would not have to define this problem). Thus, we need not consider all variants in the current path when evaluating edge weights. Instead, we could limit the number of variants to consider, from the current position in the path *i*, back some small and sensible number of steps *L*:

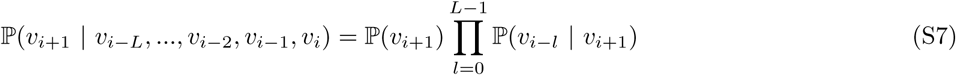

Additionally, to overcome inaccuracies encountered through floating point error when performing mathematical operations on very small decimals, **Gretel** uses log probabilities instead. Via the log identity *log*(*ab)* = *log*(*a*) + *log*(*b*) the product of the conditional probabilities becomes a sum of the log conditional probabilities:

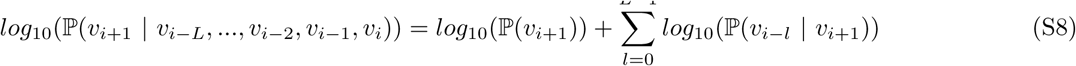

We define *L* as the the ‘lookback’ size, the number of variants of the current path to consider when selecting *v_i_*_+1_. Conveniently, there is a reasonable intuition available for selecting a value for *L*: the mean number of SNP sites covered by the observed reads. Thus we avoid the scenario of introducing an algorithmically influential but difficult to optimize parameter, such as *k*-mer size for metagenomic assembly.

### 5 Estimation of probabilities

Equation S9 provides an estimate for the marginal distribution of a symbol *β* appearing at position *j.*

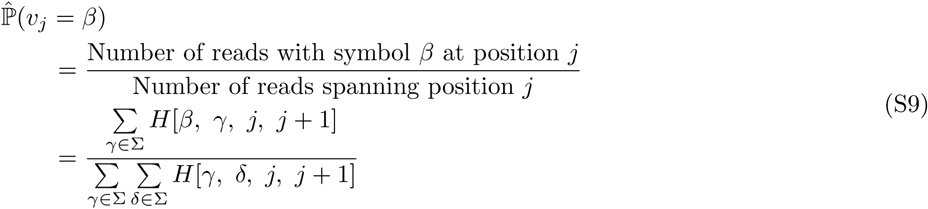

Equation S8 provides an estimate for the conditional distribution of symbol *α* appearing at position *i* given that *β* was observed at position *j*.

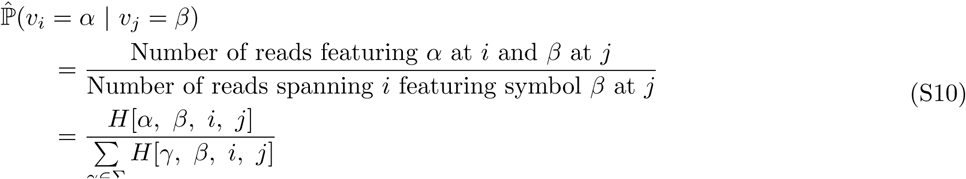

### 6 Smoothing

To avoid the potential of dividing by 0 when using Equation S10 in cases where a suitable read spanning *i* and *v*_j_ = *β* does not exist, we apply Laplace smoothing to effectively add a dummy support read. Future work will investigate alternative smoothing methodology.

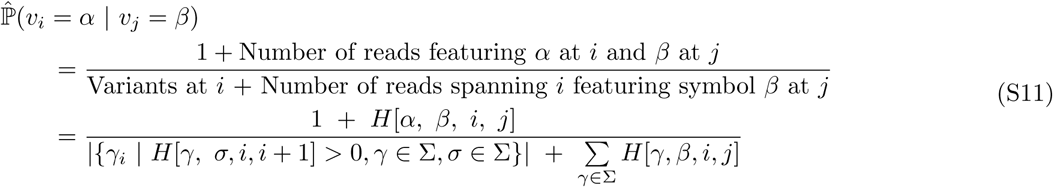

### 7 Reweighting

The paths generated by **Gretel** are probabilistic, but not stochastic. For a given *H*, **Gretel** will always return the same path. After a path has been constructed we can perform some transformation of *H* to prevent repetitive generation of the same path and return the next most likely path on the next iteration instead.

Given a path *ĥ*, we inspect the marginal distribution of each element of the path in order to find the smallest marginal. **Gretel** iterates over each element *ĥ*[*i*] in the path, and uses the **Hansel** interface to reweight the element *H*[*ĥ*[*i*],*ĥ*[*i* + 1],*i*,*i* + 1] by subtracting the result of multiplying the smallest marginal by the original value for that observation in *H*:

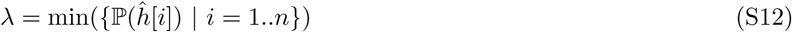

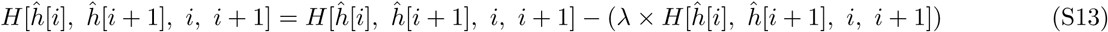

In practice *λ* is capped by **Gretel** in an attempt to stop aggressive reweighting that might otherwise prevent the recovery of closely related haplotypes.

### 8 Stopping criterion

After multiple iterations of path finding and subsequent reweighting, elements in *H* will begin to approach 0, causing edges in the graph to become unavailable for traversal. **Gretel** will immediately terminate upon encountering a node in the graph with no viable outgoing edges. That is, the selected symbol at the current i has no non-zero weighted edges to traverse between SNP positions (*i*, *i* + 1) in the graph. Alternatively, if this criterion is not reached after 100 iterations (haplotypes), **Gretel** aborts.

### 9 Haplotype scoring

**Gretel** can score and rank the haplotypes it recovers. For a completed haplotype, *ĥ*, we compute its likelihood based upon the sum of the marginal log probabilities for each element of *ĥ* given the current state of *H*.

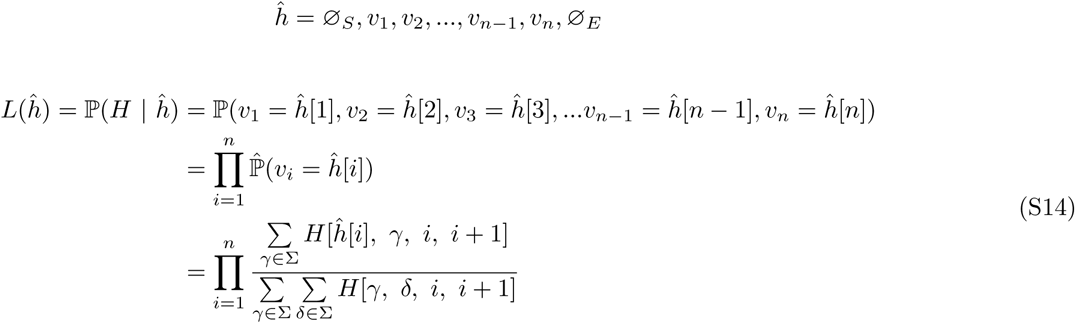

To overcome the potential for floating point arithmetic error (Equation S8), we calculate and report the log likelihood.

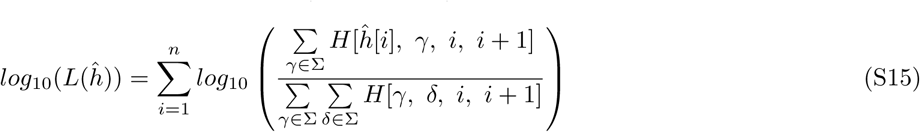

### 10 Metahaplomes from real genes (DHFR): Additional plots

**Figure 1:**
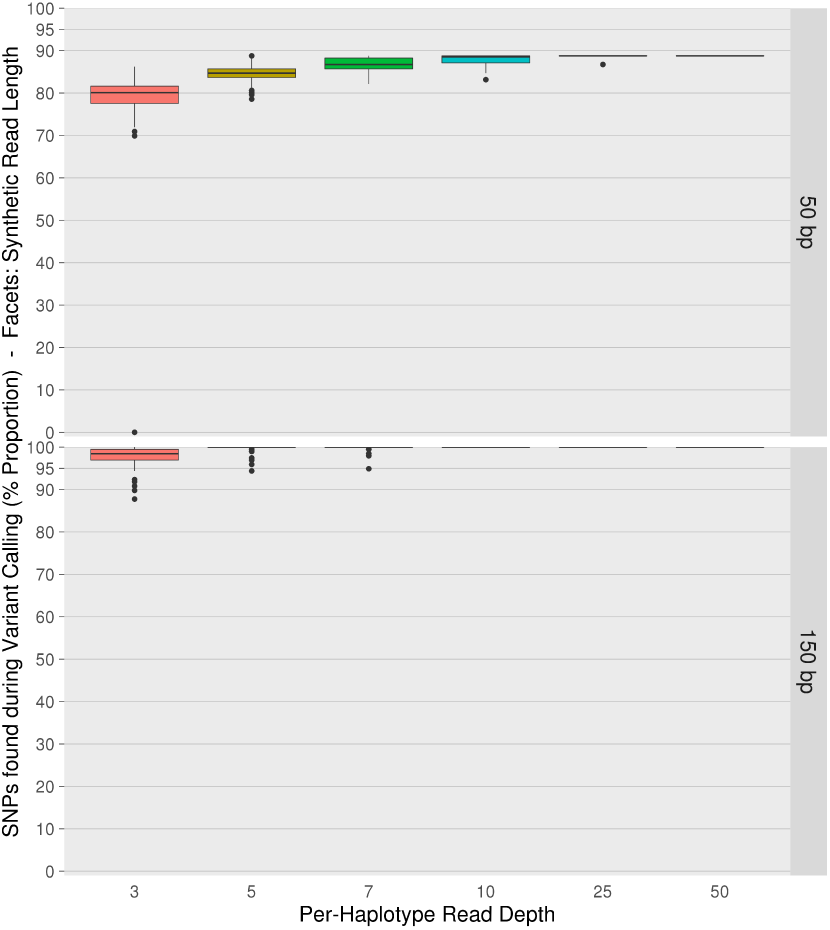
Proportion of the 196 SNPs across the five *DHFR* haplotypes that were correctly discovered by our snpper tool (y-axes) during variant calling of each of the synthetic read sets. Read sets are split by per-haplotype read depth (x-axis) and read length (row facets), each box-with-whiskers summarises 100 read sets. Uncalled SNPs prevent full recovery of one or more of the input haplotypes.

**Figure 2:**
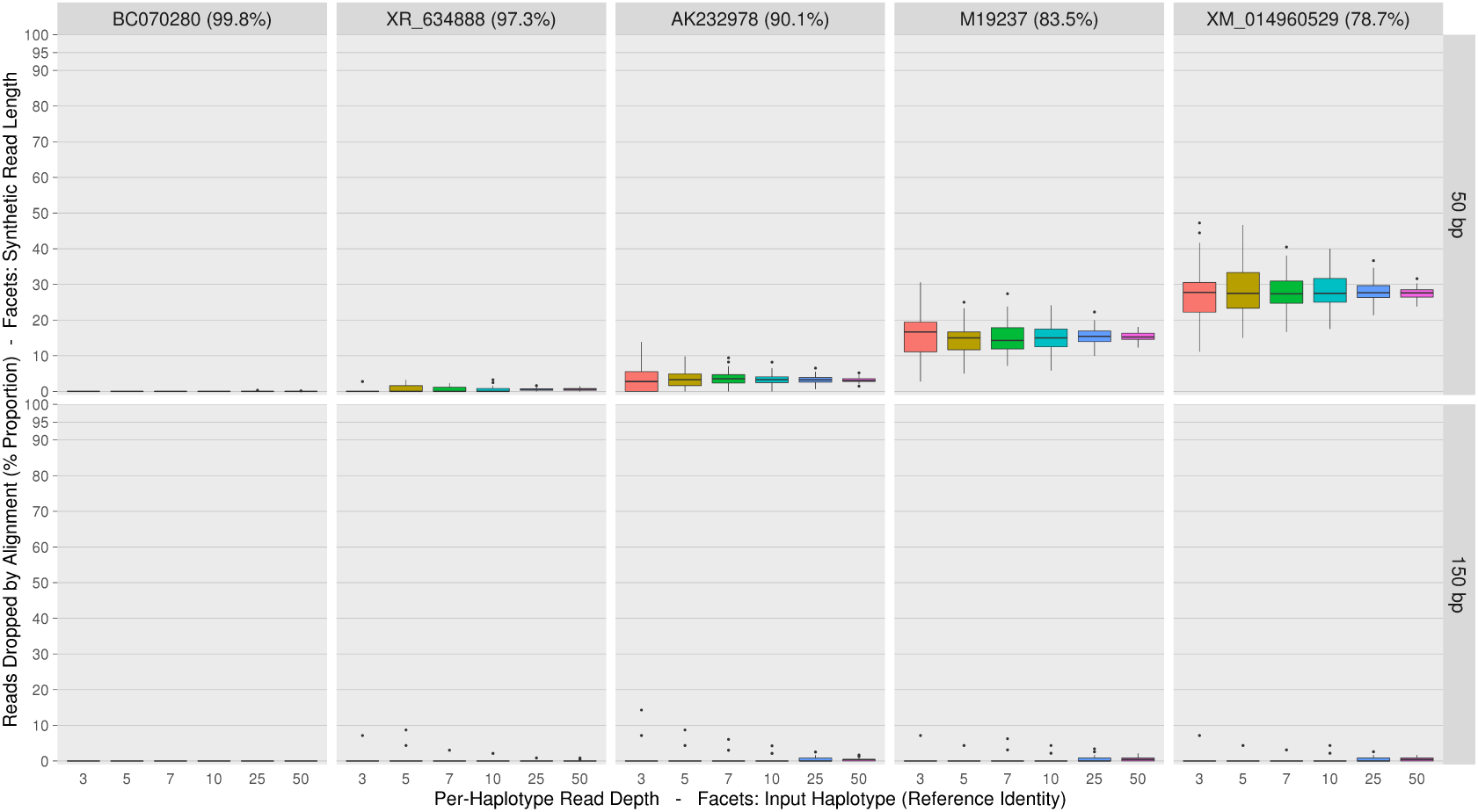
Proportion of synthetic reads from read sets dropped (y-axes) by the chosen aligner bowtie2 (see Listing 1 for parameters used), for each input *DHFR* gene (column facets). Read sets are split by per-haplotype read depth (x-axis) and read length (row facets), each box-with-whiskers summarises the proportion of a particular gene’s synthetic reads dropped, across 100 read sets. Dropped reads can impact the calling of SNPs and reduces the available evidence for the recovery of those haplotypes.

### 11 Metahaplomes with many haplotypes: Influenza A Segment 7

In order to test our methods on a data set with many haplotypes and many variants we obtained 446 Influenza A Segment 7 sequences from GenBank, requesting any sequences deposited during December 2016. After removing duplicates, and sequences with 99.0% or greater identity to any other candidates, 60 remained.

These 60 remaining sequences had varying lengths. 25 of the 60 sequences were of one particular length: 982 bp. This provided a suitable selection criteria with which to construct our metahaplome (Table 1). From the 25, *KP404423.1* was selected at random to act as the pseudo-reference for alignment. The pool of 24 remaining sequences act as haplotypes from which we generated synthetic reads via the same method used for our simulations and DHFR metahaplomes. Variants were called in the same way as our previous experiments, determining any site without a unanimous consensus as a variant.

**Table 1:** Accession numbers for the 25 Influenza A Segment 7 sequences used in our analysis. The randomly selected pseudo-reference is marked with an asterisk.

|  |  |  |  |  |
| --- | --- | --- | --- | --- |
| *KP404183.1* | *KP404287.1* | *KX215927.1* | *KY171305.1* | *KY272977.1* |
| *KP404207.1* | *KP404343.1* | *KY170930.1* | *KY171393.1* | *KY327641.1* |
| *KP404247.1* | *KP404383.1* | *KY171026.1* | *KY171401.1* | *KY327765.1* |
| *KP404255.1* | *KP404423.1\** | *KY171242.1* | *KY171489.1* | *KY327767.1* |
| *KP404271.1* | *KT717277.1* | *KY171257.1* | *KY171507.1* | *KY327784.1* |

\*Pseudo-reference

We followed the same experimental design as per the *DHFR* data sets, yielding 1,200 sets of reads (2 read sizes, 6 per-haplotype coverage levels, 100 replicates), each containing synthetic reads from the 24 haplotypes. A multiple sequence alignment of the 24 haplotypes determined the number of true SNPs to be 277. The mean number of called SNPs across all read sets was 276.02.

**Gretel** was used on each read set, returning 1,200 sets of recovered haplotypes. Each set of output haplotypes is compared pairwise against the input 24 haplotypes, calculating the Hamming distance at sites determined to be heterogeneous via a multiple sequence alignment. We report the proportion of correctly recovered heterogeneous sites, as a percentage.

Figure 3 aggregates the proportions of SNPs that were correctly recovered by **Gretel** over the 24 haplotypes under varying conditions of read size and depth. At this level of variation we produce high accuracy recoveries with reads of 50 bp. Read sets with at least 7x per-haplotype depth and 150 bp length make recoveries across the 24 haplotypes with an average accuracy of 92%. Notably, increasing read length and coverage does little to improve recovery accuracy, even at 250 bp (not shown). There appears to be an upper-limit on **Gretel**’s performance on this particular data set. As discussed in our *DHFR* Results, this could be due to dropped reads obscuring the existence of SNPs, and denying evidence of the variation that exists on that haplotype. Additionally, the shared variation in such a closely related set of sequences can cause paths in the true metahaplome to converge. Our data suggests that even in a metahaplome with 24 haplotypes, it is possible to make accurate recoveries (>90%) with reads of 50 bp.

**Figure 3:**
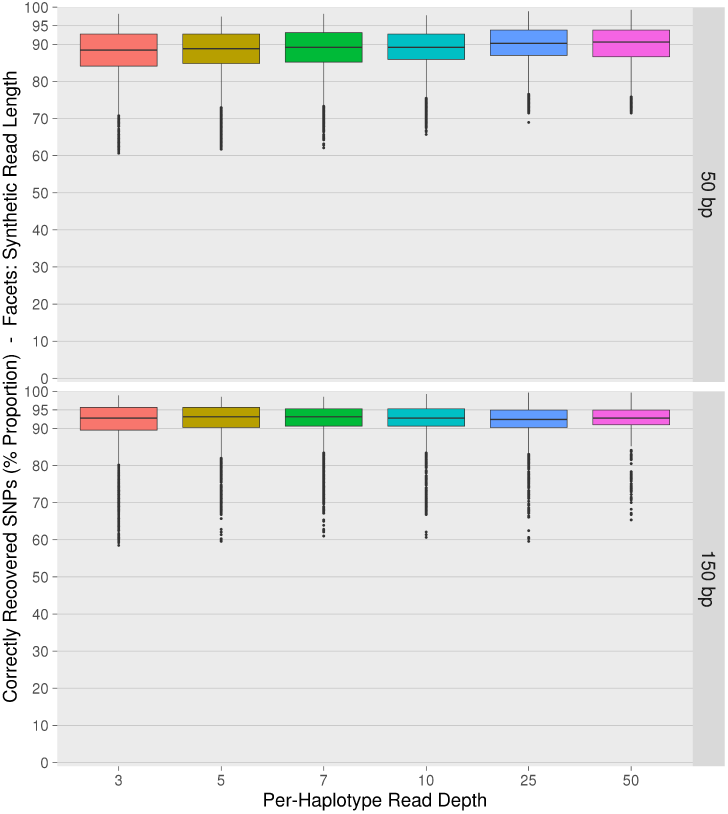
Boxplots aggregating the proportion of variants correctly recovered (y-axes) for the 24 input Influenza A Segment 7 sequences by **Gretel**. We generated reads from the *Flu-A7* metahaplome at 6 different per-haplotype read depths (x-axes) between 3 and 50x, 2 read lengths (50 bp and 150 bp row facets) with 100 replicates. Individual box-with-whiskers describe the recovery rates across the 24 haplotypes, from reads of a per-haplotype depth and size, over the 100 replicates. There appears to be an upper limit on the recoveries that are possible from this data set.

## References

[1] G. Vernikos, D. Medini, D. R. Riley, and H. Tettelin. Ten years of pan-genome analyses. Current Opinion Microbiology, 23:148–54, 2015.

[2] Richard A. Gibbs, John W. Belmont, Paul Hardenbol, Thomas D. Willis, Fuli Yu, Huanming Yang, Lan-Yang Ch’ang, Wei Huang, Bin Liu, Yan Shen, et al. The international HapMap project. Nature, 426(6968):789–796, 2003.

[3] R. Andino and E. Domingo. Viral quasispecies. Virology, 479–480:46—51, 2015.

[4] Federico M Lauro, Diane McDougald, Torsten Thomas, Timothy J Williams, Suhelen Egan, Scott Rice, Matthew Z DeMaere, Lily Ting, Haluk Ertan, Justin Johnson, et al. The genomic basis of trophic strategy in marine bacteria. Proceedings of the National Academy of Sciences, 106(37):15527–15533, 2009.

[5] Sharon A Huws, Joan E Edwards, Christopher J Creevey, Pauline Rees Stevens, Wanchang Lin, Susan E Girdwood, Justin A Pachebat, and Alison H Kingston-Smith. Temporal dynamics of the metabolically active rumen bacteria colonizing fresh perennial ryegrass. FEMS Microbiology Ecology, 92(1):fiv137, 2016.

[6] Francesco Rubino, Ciara Carberry, Sinead M Waters, David Kenny, Matthew S McCabe, and Christopher J Creevey. Divergent functional isoforms drive niche specialisation for nutrient acquisition and use in rumen microbiome. The ISME Journal, 2017.

[7] J. Jung, L. Philippot, and W. Park. Metagenomic and functional analyses of the consequences of reduction of bacterial diversity on soil functions and bioremediation in diesel-contaminated microcosms. Nature Scientific Reports, 6:23012, 2016.

[8] Chen Zhang and Se-Kwon Kim. Research and application of marine microbial enzymes: status and prospects. Marine Drugs, 8(6):1920–34, 2010.

[9] V. Kuleshov, C. Jiang, W. Zhou, F. Jahanbani, S. Batzoglou, and M. Snyder. Synthetic long-read sequencing reveals intraspecies diversity in the human microbiome. Nature Biotechnology, 34:64 – 69, 2016.

[10] Giuseppe Lancia, Vineet Bafna, Sorin Istrail, Ross Lippert, and Russell Schwartz. SNPs problems, complexity, and algorithms. In Algorithms—ESA 2001, pages 182–193. Springer, 2001.

[11] Rudi Cilibrasi, Leo Van Iersel, Steven Kelk, and John Tromp. On the complexity of several haplotyping problems. In Algorithms in Bioinformatics, pages 128–139. Springer, 2005.

[12] Ross Lippert, Russell Schwartz, Giuseppe Lancia, and Sorin Istrail. Algorithmic strategies for the single nucleotide polymorphism haplotype assembly problem. Briefings in Bioinformatics, 3(1):23–31, 2002.

[13] Giuseppe Lancia. Algorithmic approaches for the single individual haplotyping problem. RAIRO-Operations Research, 50(2):331–340, 2016.

[14] Filippo Geraci. A comparison of several algorithms for the single individual SNP haplotyping reconstruction problem. Bioinformatics, 26(18):2217–2225, 2010.

[15] P. Edge, V. Bafna, and V. Bansal. HapCUT2: robust and accurate haplotype assembly for diverse sequencing technologies. Genome Research, Advance access (10.1101/gr.213462.116), 2016.

[16] D. Aguiar and S. Istrail. HapCompass: a fast cycle basis algorithm for accurate haplotype assembly of sequence data. Journal of Computational Biology, 19(6):577–590, 2012.

[17] Ehsan Motazedi, Richard Finkers, Chris Maliepaard, and Dick de Ridder. Exploiting next-generation sequencing to solve the haplotyping puzzle in polyploids: a simulation study. Briefings in Bioinformatics, page bbw126, 2017.

[18] Derek Aguiar and Sorin Istrail. Haplotype assembly in polyploid genomes and identical by descent shared tracts. Bioinformatics, 29(13):i352–i360, 2013.

[19] T. Namiki, T. Hachiya, H. Tanaka, and Y. Sakakibara. MetaVelvet: An extension of Velvet assembler to de novo metagenome assembly from short sequence reads. Nucleic Acids Res, 40(20):e155, 2012.

[20] Sébastien Boisvert, Frédéric Raymond, Élénie Godzaridis, François Laviolette, and Jacques Corbeil. Ray Meta: scalable de novo metagenome assembly and profiling. Genome Biology, 13(12):R122, 2012.

[21] John Vollmers, Sandra Wiegand, and Anne-Kristin Kaster. Comparing and evaluating metagenome assembly tools from a microbiologist’s perspective-not only size matters! PLoS One, 12(1):e0169662, 2017.

[22] D. Y. C. Brandt, V. R. C. Aguiar, B. D. Bitarello, K. Nunes, J. Goudet, and D. Meyer. Mapping bias overestimates reference allele frequencies at the HLA genes in the 1000 Genomes Project Phase I Data. G3: Genes – Genomes – Genetics, 5(5):931–941,2015.

[23] Erik Garrison et al. vg: the variation graph toolkit. https://github.com/vgteam/vg, 2016.

[24] Andrew Rambaut and Nicholas C. Grass. Seq-Gen: an application for the Monte Carlo simulation of DNA sequence evolution along phylogenetic trees. Computer applications in the biosciences: CABIOS, 13(3):235–238, 1997.

[25] Siegfried Schloissnig, Manimozhiyan Arumugam, Shinichi Sunagawa, Makedonka Mitreva, Julien Tap, Ana Zhu, Alison Waller, Daniel R Mende, Jens Roat Kultima, John Martin, et al. Genomic variation landscape of the human gut microbiome. Nature, 493(7430):45–50, 2013.

[26] Moni Sharma and Prem MS Chauhan. Dihydrofolate reductase as a therapeutic target for infectious diseases: opportunities and challenges. Future Medicinal Chemistry, 4(10):1335–1365, 2012.

[27] Francesca Di Giallonardo, Armin Töpfer, Melanie Rey, Sandhya Prabhakaran, Yannick Duport, Christine Leemann, Stefan Schmutz, Nottania K. Campbell, Beda Joos, Maria Rita Lecca, Andrea Patrignani, Martin Däumer, Christian Beisel, Peter Rusert, Alexandra Trkola, Huldrych F. Günthard, Volker Roth, Niko Beerenwinkel, and Karin J. Metzner. Full-length haplotype reconstruction to infer the structure of heterogeneous virus populations. Nucleic Acids Res, 42(14):e115, 2014.

[28] J.M. Cuevas, R. Geller, R. Garijo, J. López-Aldeguer, and R. Sanjuán. Extremely high mutation rate of HIV-1 in vivo. PLoS Biol, 13(9):e1002251, 2015.

[29] NCBI Resource Coordinators. Database resources of the National Center for Biotechnology Information. Nucleic Acids Research, 44(Database issue):D7–D19, 2016.

[30] Liang Shi, Hailiang Dong, Gemma Reguera, Haluk Beyenal, Anhuai Lu, Juan Liu, Han-Qing Yu, and James K Fredrickson. Extracellular electron transfer mechanisms between microorganisms and minerals. Nature Reviews Microbiology, 14(10):651–662, 2016.

[31] Priscilla EM Purnick and Ron Weiss. The second wave of synthetic biology: from modules to systems. Nature Reviews Molecular Cell Biology, 10(6):410–422, 2009.

[32] Karel Břinda, Valentina Boeva, and Gregory Kucherov. OCOCO: the first online consensus caller. https://github.com/karel-brinda/ococo, 2016.

[33] Soyeon Ahn and Haris Vikalo. Joint haplotype assembly and genotype calling via sequential Monte Carlo algorithm. BMC Bioinformatics, 16(1):223, 2015.

[34] D. Aguiar. HapCompass manual. Technical report, Brown University, 2014.

[35] Rebecca Rose, Bede Constantinides, Avraam Tapinos, David L Robertson, and Mattia Prosperi. Challenges in the analysis of viral metagenomes. Virus Evolution, 2(2):vew022, 2016.

[36] Niko Beerenwinkel, Huldrych F Günthard, Volker Roth, and Karin J Metzner. Challenges and opportunities in estimating viral genetic diversity from next-generation sequencing data. Frontiers in Microbiology, 3:329, 2012.

[37] Armin Töpfer, Osvaldo Zagordi, Sandhya Prabhakaran, Volker Roth, Eran Halperin, and Niko Beerenwinkel. Probabilistic inference of viral quasispecies subject to recombination. Journal of Computational Biology, 20(2):113–123, 2013.

[38] S. Prabhakaran, M. Rey, O. Zagordi, N. Beerenwinkel, and V. Roth. HIV haplotype inference using a propagating dirichlet process mixture model. IEEE/ACM Trans. Comput. Biol. Bioinform., pages 182–191, 2013.

[39] Aridaman Pandit and Rob J de Boer. Reliable reconstruction of HIV-1 whole genome haplotypes reveals clonal interference and genetic hitchhiking among immune escape variants. Retrovirology, 11(1):56, 2014.

[40] Duleepa Jayasundara, Isaam Saeed, Suhinthan Maheswararajah, B. C. Chang, S.-L. Tang, and Saman K Halgamuge. ViQuaS: an improved reconstruction pipeline for viral quasispecies spectra generated by next-generation sequencing. Bioinformatics, 31(6):886–96, 2015.

[41] V. Kuleshov. Probabilistic single-individual haplotyping. Bioinformatics, 30(17):i379–85, 2014.

[42] Alessandro Panconesi and Mauro Sozio. Fast hare: A fast heuristic for single individual SNP haplotype reconstruction. In International Workshop on Algorithms in Bioinformatics, pages 266–277. Springer, 2004.

[43] Hirotugu Akaike. Likelihood of a model and information criteria. Journal of Econometrics, 16(1):3–14, 1981.

[44] Matthew B. Scholz, Chien-Chi Lo, and Patrick S. G. Chain. Next generation sequencing and bioinformatic bottlenecks: the current state of metagenomic data analysis. Current Opinion in Biotechnology, 23(1):9–15, 2012.

[45] Zamin Iqbal, Mario Caccamo, Isaac Turner, Paul Flicek, and Gil McVean. De novo assembly and genotyping of variants using colored de Bruijn graphs. Nature Genetics, 44(2):226–232, 2012.

[46] D. R. Zerbino and E. Birney. Velvet: algorithms for de novo short read assembly using de Bruijn graphs. Genome Research, 18:821–829, 2008.

[47] B. Langmead and S. Salzberg. Fast gapped-read alignment with Bowtie 2. Nature Methods, 9:357–359, 2012.

[48] H. Li, B. Handsaker, A. Wysoker, T. Fennell, J. Ruan, N. Homer, G. Marth, G. Abecasis, R. Durbin, and 1000 Genome Project Data Processing Subgroup. The sequence alignment/map (SAM) format and SAMtools. Bioinformatics, 25:2078–9, 2009.

[49] M. DePristo, E. Banks, R. Poplin, K. Garimella, J. Maguire, C. Hartl, A. Philippakis, G. del Angel, M. A. Rivas, M. Hanna, A. McKenna, T. Fennell, A. Kernytsky, A. Sivachenko, K. Cibul-skis, S. Gabriel, D. Altshuler, and M. Daly. A framework for variation discovery and genotyping using next-generation DNA sequencing data. Nature Genetics, 43:491–498, 2011.

[50] R. C. Edgar. MUSCLE: a multiple sequence alignment method with reduced time and space complexity. BMC Bioinformatics, 5(1):113, 2004.

[51] Bette Korber, Brian T Foley, C Kuiken, Satish K Pillai, Joseph G Sodroski, et al. Numbering positions in HIV relative to HXB2CG. Human retroviruses and AIDS, 3:102–111, 1998.

